# TEAD Palmitoylation Inhibitors are Brain Penetrant and Enhance Survival in Preclinical Models of Recurrent Glioblastoma in Combination with Standard-of-Care Therapy

**DOI:** 10.64898/2026.11.18.767633

**Authors:** Thenzing J. Silva-Hurtado, Balagopal Pai, Mohammad Imran K. Khan, Debdatta Halder, Anne Marie Barrette, Timothy E. Richardson, Anirudh Sattiraju, Roland H. Friedel, Hongyan Zou, Dolores Hambardzumyan, Raymund L. Yong, Tracy T. Tang, Nadejda M. Tsankova

## Abstract

**Purpose:** Oncogenic addiction to YAP-TEAD signaling drives adaptive plasticity, therapeutic resistance, invasion and tumor progression in many cancers, including in glioblastoma (GBM), and represents an actionable vulnerability to explore as adjuvant therapy during tumor recurrence. This study evaluated the translational potential of orally bioavailable Vivace Therapeutics (VT) TEAD palmitoylation inhibitors in preclinical GBM models.

**Experimental Design:** The effects of VT103 and VT104 on GBM proliferation, migration, and TEAD activity were first assessed across patient-derived GBM cell lines with diverse genomic drivers. The drugs’ pharmacokinetics (PK) and biodistribution were measured using mass spectrometry in mice bearing patient-derived orthotopic xenograft (PDX) GBM. Therapeutic efficacy was evaluated in two genetically distinct PDX models of recurrent GBM, administering VT104 as monotherapy or in combination with standard-of-care radiotherapy plus temozolomide (RT+TMZ).

**Results:** VT104/VT103 treatment significantly impaired GBM proliferation and migration in- vitro, with concomitant downregulation of TEAD-target expression. PK studies confirmed sustained VT104 plasma levels and robust brain biodistribution in PDX mice, with brain:plasma ratio >150% at therapeutic dosing without systemic toxicity. While VT104 monotherapy showed modest anti-tumor effects, combination therapy (VT104+RT+TMZ) significantly extended overall survival in both recurrent GBM PDX models. The survival benefit was most prominent in *EGFR*-mutant GBM and corresponded to significant decrease in overall and infiltrative tumor burden. Mechanistically, VT104 potentiated TMZ-dependent γ-H2AX genotoxic stress and RT+TMZ synergized with VT104 to abolish TEAD activity.

**Conclusion:** Integrating YAP-TEAD inhibition with standard therapy introduces a clinically- relevant paradigm for synergistic treatment in recurrent GBM that both inhibits tumor infiltration and reduces therapeutic resistance to chemoradiation.

## Introduction

Isocitrate dehydrogenase (IDH)-wildtype glioblastoma (GBM) is the most prevalent and aggressive primary brain neoplasm in adults, characterized by rapid cellular proliferation, diffuse parenchymal infiltration, and profound resistance to therapy ^1–4^. The current standard of care treatment in patients with GBM, comprising maximal safe surgical resection followed by radiotherapy (RT) and adjuvant temozolomide (TMZ) chemotherapy ^5^, remains palliative. Most patients ultimately succumb to recurrence within a median progression free survival of only 6-9 months and a dismal median overall survival of 12-18 months ^6^. The management of recurrent GBM represents one of the most formidable challenges in neuro-oncology, as second line interventions, including surgical re-resection, systemic chemotherapy, re-irradiation and targeted agents have consistently demonstrated modest outcomes without clear benefit in overall survival ^7, 8^. This persistent therapeutic plateau exposes a gap in effective treatment options and an urgent need for novel strategies that target the molecular determinants of adaptative resistance leading to tumor recurrence ^9^.

In recent years, the Hippo signaling pathway has emerged as a critical regulator of adaptive plasticity and therapeutic resistance in cancer ^10, 11^, including in GBM ^12, 13^. Dysregulation of this pathway results in aberrant activation of the transcriptional co-activators YAP and TAZ ^14^, which drive TEAD-mediated transcriptional programs that promote GBM tumor growth, proliferation, and tumor invasion ^15–18^. Crosstalk between Hippo YAP-TEAD and other hyperactivated oncogenic signaling inputs, including receptor tyrosine kinase (RTK) pathways, reinforce its role as a central node of adaptive resistance in GBM and other cancers ^10–12^.

Pharmacologic disruption of the YAP/TAZ–TEAD transcriptional complex has consequently emerged as a rational therapeutic strategy ^12^. Several prior studies have shown efficacy for the YAP-TEAD inhibitor Verteporfin in GBM, both in-vitro and in-vivo ^15, 16, 19–21^. However, Verteporfin is known to have activity outside of YAP-TEAD signaling, shows limited brain penetrance and a short half-life, and is not amenable to oral administration due to its lipophilic structure ^22^, thus limiting its translational potential. Newer small-molecule TEAD palmitoylation inhibitors with superior pharmacokinetic / pharmacodynamic (PK/PD) profiles have recently emerged ^23, 24^. TEAD palmitoylation inhibitors target the conserved hydrophobic pocket required for TEAD auto- palmitoylation, destabilizing TEAD and preventing its interaction with YAP/TAZ ^24^. Among these, the class of inhibitors developed by Vivace Therapeutics (VT) selectively block TEAD palmitoylation, suppressing oncogenic transcriptional programs that sustain tumor growth and survival, and are available as an oral formulation pill. VT compounds were discovered through a rational screen for potent YAP-TEAD inhibitors, based on reporter selection that minimizes off target action ^25^. Preclinical studies have demonstrated favorable PK profiles of VT compounds outside of the central nervous system (CNS), including high oral bioavailability (>75%), long half- life (>12h), and dose-dependent tumor regression with minimal systemic toxicity ^25–28^. Indeed, VT compounds have shown anti-tumor efficacy in mesotheliomas, meningiomas, and schwannomas ^25–27^ and the VT clinical candidate has demonstrated clinical response and is well tolerated in ongoing clinical trials for these tumors ^28^.

Despite their promising preclinical and clinical profile, the role of VT agents in primary brain tumors remains largely unknown. Here, we investigate the PK/PD properties of VT103 and VT104 TEAD palmitoylation inhibitors ^25^, specifically in the CNS, and evaluate VT104’s therapeutic efficacy in patient-derived xenograft (PDX) models of recurrent GBM, alone or in combination with RT and TMZ. By targeting YAP-TEAD dependent transcriptional programs that maintain glioma stem cell plasticity and therapeutic resistance, this study aims to define the translational relevance of TEAD palmitoylation VT inhibitors in recurrent GBM.

## Material and methods

### Inclusion criteria

In-vitro functional studies included patient-derived cell lines from both female and male sex. In- vivo studies were performed with adult female mice in order to sex-match the patient-derived cell line donor with the host recipient.

### Proliferation and migration in-vitro assays

Functional tumor properties were assessed in low-passage patient-derived GBM cell lines with distinct genomic drivers: G-13063 with *PDGFRA* alteration; G-12746 with *EGFR* alteration; and G-16302 with *NF1* alteration ^16, 17^. Cells were cultured on laminin-coated plates in serum-free neurosphere medium, as previously described ^17, 29^. Viability studies for IC_50_ dose were performed using the CellTiter Glo assay (Promega, G7570). Cell proliferation after vehicle or drug treatment (IC_50_ dose) was assessed using the countess 750 automated cell counter (Thermo Fisher, C10228, RRID:SCR_025370). Briefly, a total of 25,000 GBM cells were seeded onto 10 μg/ml laminin-coated 12-well plates (Corning, 40232) and cultured in serum-free neurosphere medium with growth factors. Cells were harvested at 0, 24, 48, and 72h, and viable cell numbers were quantified to generate a growth curve. For spheroid formation, 2500 viable cells were seeded onto 1% agarose-coated wells and grown to spheroids for at least 48h. Individual spheroids of similar size were then transferred to laminin-precoated, round bottom 96-well plates (Nunc, 268200) and treated with drug (IC_50_ dose) or vehicle, in cell culture medium that lacks growth factors. Area of confluent spheroid cell dispersion was measured at baseline (0h), at 24h, and at 36h after treatment, under bright field microscopy using ImageJ (RRID:SCR_003070).

### Recurrent GBM orthotopic xenograft mouse modeling

Orthotopic xenografts were generated in 2-month-old immunocompromised patient sex- matched (female) mice, IcrTac:ICR-Prkdcscid strain (Taconic, RRID:IMSR_TAC:ICRSC) with lymphocyte deficiency but retained microglial activity. Two diffusely infiltrative patient-derived recurrent GBM cell lines were used: G-16302 (*NF1*-mutant) and G-12315 (*EGFR* amplified + *EGFR vIII* mutant)^30^, maintained and expanded under stem cell / tumor-initiating conditions following established protocols ^16, 17, 31, 32^. Anaesthetized mice were orthotopically implanted with viable G-16302 or G-12315 GBM cells (2.0x10^5^ cells) in 2μl Opti-MEM medium (Gibco, 31985070) into the striatum using the following coordinates: 2.0mm right lateral to bregma and 3.0mm deep ^16^. Intracranial delivery was performed using a 10ul microsyringe (Hamilton, 80030) mounted on a manual stereotaxic frame (Stoelting, 51500), at a rate of 0.5ul /min with gradual withdrawal. To ensure optimal cell viability and minimize experimental variability, surgeries were limited to a maximum of five mice per day. Animals were monitored continuously until recovery from anesthesia, placed in a heated recovery chamber, and returned to their home cages upon regaining normal mobility.

### VT brain pharmacokinetic / pharmacodynamic studies

For pharmacokinetics studies, drug levels of VT103 and VT104 (Vivace Therapeutics) were measured in plasma and perfused brain tissue of tumor-bearing G-16302 PDX mice, treated at 10 mg/kg daily for two weeks and every other day for five additional weeks via oral gavage, using high-performance liquid chromatography with mass spectrometry (LC-MS/MS) at Quintara Discovery, Hayward California. Mice were euthanized in a staggered manner, 4h and 24h after the final drug administration. Blood (at least 200 µL) was collected via cardiac puncture in prechilled EDTA-coated tubes (BD microtainer, 365974), centrifuged at 8,000 rpm for 5 min at 4 °C, and the plasma fraction (∼100 µL) was carefully isolated and snap-frozen on dry ice and stored at –80 °C until analysis. To minimize blood contamination in tissue drug quantification, mice were transcardially perfused with phosphate-buffered saline (1XPBS pH7.2 Gibco, 20012- 027) for 3 minutes, using a peristaltic pump (Fisher Scientific, 15357547). Following perfusion, organs were harvested individually, blotted and weighed using a high-sensitivity analytical balance. Tissues were snap-frozen on dry ice and stored at –80 °C until analysis. For LC-MS/MS analysis, brain tissue was homogenized with 2 volumes of 20% acetonitrile in water and further diluted two-fold with blank mouse plasma. A 20 µL aliquot of plasma or plasma-diluted brain homogenate was extracted with 100 µL of 90% acetonitrile in water with 5 mM ammonium acetate and 0.1% acetic acid containing an internal standard (Terfenadine). The mixture was vortexed on a shaker for 15 minutes and subsequently centrifuged at 3220 g for 15 minutes. A 70 µL aliquot of the supernatant was mixed with 70 µL of 70% acetonitrile in water for the injection to the LC/MS/MS using a C18 50 x 2.1 mm, 1.7 µm column at 40 °C and 0.5mL/min flow rate. Calibration standards and quality control samples were prepared by spiking the test compounds into blank mouse plasma and processed in parallel with the unknown samples in the same batch. For pharmacodynamics studies, drug-treated gliomaspheres (IC_50_ dose VT103 and VT104, 72h after treatment) and G-16302 / G-12315 PDX harvested tissues were used to quantify expression of TEAD-target genes using quantitative (q)RT-PCR (Applied Biosystems, QuantStudio 5, RRID:SCR_020240). Briefly, RNA was extracted using TRIzol (Invitrogen, 15596026), followed by chloroform (Fisher Scientific, 15498679) phase separation, isopropanol (Fisher Scientific, 327270010) precipitation, 75% alcohol wash, and resuspension in nuclease-free water (Invitrogen Ultrapure Distilled Water, 11538646). RNA concentration was assessed using a Nanodrop spectrophotometer (Nanodrop 1000 Thermo Scientific, RRID:SCR_016517). RNA was transcribed into complementary DNA (cDNA) and qRT-PCR was performed using SYBR Green Master Mix (Fisher Scientific, A46109) on Real-Time PCR System. TEAD-target gene expression levels (*CTGF*, *CYR61, EGFR)* were calculated using the 2^-ΔΔCT^ method, normalized to housekeeping gene (*Rpl13a, RPLPO,* or *18S)* and relative to vehicle control. All reactions were performed in technical triplicates using custom-designed primer sequences specific to either human or mouse genome (see Supplementary Table 2).

### Drug and radiation treatments

For animal studies, VT104 was administered via oral gavage (Instech, FTP-20gax38mm) at 10mg/kg/day, formulated in 5% DMSO (Sigma Aldrich,D2650), 10% Solutol (MedChemExpress, HY-Y1893), and 85% D5W, where D5W is 5% glucose (Fisher Scientific, D16-500). Treatment began 2 weeks after tumor implantation and continued for 7 weeks (for IVIS and PK studies), until end-survival point (for survival and histology studies), or for 3 weeks (for validation in-vivo G- 12315 studies). VT104 was administered daily for the first 2-3 weeks, and then every other day. RT+TMZ treatment began 4 weeks after tumor implantation and consisted of concomitant TMZ (25mg/kg/day via oral gavage, MedChemExpress, HY-17364) and RT (2Gy/day) administered for 10 days (five days on, two days off), using standard preclinical dosing as before ^33^. Focal cranial irradiation was performed using an X-ray biological irradiator (RS 2000 Biological Irradiator, Rad Source Technologies Inc). Anaesthetized mice were placed in a mouse irradiation chamber (shelf position 1) exposing only the cranium to radiation, with the rest of the body being shielded by lead protectors. Irradiation was delivered at 160 kV and 25 mA for a total exposure time of 1 minute and 40 seconds, corresponding to a 2Gy fraction. DMSO formulation (5% DMSO, 10% Solutol, and 85% D5W) was administered as a vehicle control to match VT104 treatment schedule. Mice were monitored for at least 10 minutes post-administration to ensure recovery and proper physiological response. Throughout the study, gavage was performed at the same time each day to reduce circadian variability and stress-induced differences in pharmacokinetics or behavior. Animals were returned to their home cages with *ad libitum* access to food and water after treatment.

### Survival studies

Survival efficacy was assessed in two sex-matched, orthotopic PDX models of recurrent GBM: G- 16302 and G-12315. Following tumor implantation, mice were randomly assigned to one of four different experimental groups (sample size n=10 per group): vehicle control, VT104, RT+TMZ, or VT104+RT+TMZ. Mice were monitored daily for overall health and the onset of neurological deficits. Survival was defined as the time from cell implantation to humane endpoint, determined according to institutional animal care guidelines (20% loss of peak body weight or presence of severe neurological signs such as lethargy, ataxia or marked kyphosis). Euthanasia was performed via cervical dislocation to ensure rapid death and optimal preservation of brain tissue for experimental analysis. Regarding experimental attrition, a total of 2 animals were excluded from the experiments due to low baseline body weight.

### Analyses of tumor growth and infiltration in-vivo

In-vivo bioluminescence imaging (BLI) was performed to monitor tumor growth / progression in live mice with orthotopic PDX, generated using the G-16302_akaluc patient-derived recurrent GBM cell line, genetically modified to express the bioluminescent reporter Akaluc ^32^. The In Vivo Imaging System (IVIS) Spectrum (PerkinElmer/Revvity, Waltham, MA, USA) was used for BLI, equipped with an Andor ikon CCD camera (DZ436). Images were collected and quantified using the Living Image Software (v4.7.4;PerkinElmer). Briefly, mice were injected intraperitoneally with AkaLumine-HCl luminescence substrate (Sigma-Aldrich, 808350) at a dose of 25 mg/kg as described ^32^. Luminescent images were acquired with a 5-second exposure time, binning factor of 8, f/stop 1, and a field of view of 22.6cm. Photographic images were captured with an exposure time of 0.2s, binning factor of 4 and f/stop 8. BLI measurements were performed at weekly intervals, with signal measured before substrate administration (time 0, baseline) and then at 5- minute intervals after substrate administration, for up to 30 minutes. Maximum BLI signal measured for each mouse and each week was then normalized to time 0 / baseline BLI signal. For histological analysis of tumor burden, following euthanasia, brains from survival PDX studies were carefully removed during necropsy, fixed in 4% paraformaldehyde (Fisher Scientific, AAJ19943K2) for 24 hours, dehydrated through graded alcohols, and embedded in paraffin. Coronal sections (5 µm thick) were then stained with hematoxylin and eosin (H&E) for morphological evaluation of overall and infiltrative tumor burden. Tumor area was measured using QuPath (RRID:SCR_018257) with tumor density greater than 80% annotated as tumor core and lower tumor density annotated as infiltrative edge ^16,34^. In addition, brain samples intended for molecular analysis were snap-frozen in liquid nitrogen and stored at -80 °C.

### Assessment of Genotoxic Stress

G-12315 cells were cultured and treated with VT104 (20 μM), TMZ (50 μM), IR (2Gy+2Gy) or a combination of these treatments. Cells were harvested at 96h to evaluate genotoxic stress by measuring phosphorylated histone H2AX (γ-H2AX) intensity using immunofluorescence (IF). For IF staining, cells were incubated with a primary anti γ-H2AX antibody (Cell Signaling, 2577, RRID:AB_2118010, 1:400) followed by an Alexa Fluor 647-conjugated secondary antibody (Invitrogen, A32733TR, RRID:AB_2866492, 1:200). Nuclei were counterstained and mounted using DAPI fluoromount-G mounting medium (Invitrogen, 00-4959-52). Confocal images were acquired using a Leica TCS SP8 AOBS confocal laser scanning microscope (Leica Microsystems). Intensity of γ-H2AX fluorescence was quantified using ImageJ software.

### Longitudinal GLASS dataset YAP–TEAD signature analysis

Publicly available GBM RNA-seq data with EGFR amplification status were obtained from the Glioma Longitudinal Analysis Consortium (GLASS) RNA-seq data resource ^35, 36^ via cBioPortal ^37,38^. Matched primary and recurrent IDH-wildtype GBM samples were identified using patient- level identifiers, selecting only primary GBM samples without any neoadjuvant treatment and matched first GBM recurrences where the patient received both radiation and temozolomide prior to the recurrence. This yielded n=68 non-treated (primary) and n=68 matched RT+TMZ- treated (recurrent) GBM data samples, for which EGFR amplification status was also extracted. Gene-wise z-scores were re-calculated from log-normalized TPM data ( log2(TPM+1)) across the 68 pairs of matched samples. Recurrent minus primary expression change was calculated for each gene as: Δz = recurrent tumor z-score – matched primary tumor z-score and mean- centered within each patient before visualization. YAP–TEAD activity was assessed using the CORDENONSI_YAP_CONSERVED_SIGNATURE gene set from the Molecular Signatures Database (MSigDB), which contains genes upregulated by YAP or TAZ and enriched in YAP-, TAZ-, or TEAD-dependent cancer cell lines ^39^ and has been previously shown to be downregulated after TEAD inhibition in GBM ^16^. For each matched patient, a mean YAP–TEAD signature score difference was calculated as the average recurrent minus primary Δz-score across all available genes in the MSigDB dataset. Patients were ordered from lowest to highest mean YAP–TEAD signature score difference (Δz-score), and individual genes’ Δz-score was represented in the heatmap. Mean YAP–TEAD signature Δz score was displayed as a continuous viridis-scaled annotation bar on the top. A housekeeping control signature was derived from the housekeeping gene set reported ^40^, omitting ACTB from the analysis. Heatmaps were generated in R using ComplexHeatmap ^41^, with row-wise hierarchical clustering and no column clustering. Heatmap colors were fixed from –6 to +6 Δz-score. Patient-level EGFR amplification status was annotated in binary as positive if EGFR amplification was present (magenta).

### Statistical analysis

Statistical analyses and data visualization were conducted in Prism (GraphPad Software, RRID:SCR_002798). Comparisons between two groups were performed using Student’s t-test or Welch t-test (unequal variance); for experiments involving more than two groups, two-way ANOVA was used to determine statistical p-value (p) significance. Survival data was analyzed using the Kaplan–Meier method, and statistical comparisons among groups were performed using the log-rank (Mantel–Cox) test.

### Study approval

All experimental procedures were approved by the Institutional Animal Care and Use Committee (IACUC) at the Icahn School of Medicine at Mount Sinai, in accordance with the National Institutes of Health Guide for the Care and Use of Laboratory Animals (STUDY 2016-0530). Experiments were conducted by an experienced veterinarian trained in stereotactic surgery and oral gavage techniques, with particular attention to minimizing stress, ensuring reproducibility, and maintaining animal welfare.

### Data availability

Aggregated values for all individual data points in graphs and reported means are provided in the Supporting Data Values file. Published GLASS consortium RNA-seq data was downloaded through the cBioPortal, https://www.cbioportal.org/.

## Results

### VT agents impair early tumor migration and suppress TEAD-dependent transcription across GBM subtypes

To establish the functional impact of TEAD palmitoylation inhibitors on the aggressive biology of GBM, we first evaluated VT drug effects on tumor proliferation and tumor migration in-vitro, across several low-passage patient-derived GBM cell lines with diverse genomic drivers, representatives of different molecular subtypes ^16, 17^ (Supplementary Table 1). We prioritized VT analogues with improved potency and superior oral pharmacokinetics, including VT103, a TEAD1- selective inhibitor, and VT104, a pan-TEAD inhibitor ^25^. Both drugs displayed dose-dependent inhibition of cell viability with overall similar IC_50_ values determined across cell lines (Supplementary Figure S1A). Accordingly, treatment with both VT103 and VT104 led to significant decrease in GBM cell proliferation after 48 hours, compared to DMSO vehicle control (Supplementary Figure S1B). We next examined the effects of VT treatment on GBM migration using the spheroid dispersion assay, where migration is uncoupled from proliferation by removing growth factors during cell dispersion ^16^. Notably, both VT103 and VT104 agents significantly reduced confluent GBM cell migration compared to vehicle-treated control (Figure 1A-B). Reduced migration was seen at 24 and at 36 hours post-treatment, time points when cell proliferation and doubling effects were non-significant (Supplementary Figure S1B-C). In G-13063 cells, VT103 displayed more significant inhibition of spheroid migration despite showing a much smaller decrease in cell proliferation, compared to VT104, further supporting the observation that effects on migration are not due to inhibition of cell proliferation (Figure 1A, middle; Supplementary Figure S1B, right). To correlate the observed functional effects to VTs’ mechanism of drug action, we performed pharmacodynamic analysis on the same drug-treated gliomaspheres at 36 hours. Assessment of gene expression by qRT-PCR revealed significant downregulation of YAP/TAZ-TEAD transcriptional targets, *CYR61 and CTGF*, following treatment with VT103, VT104, and Verteporfin, compared to vehicle control (Figure 1C, Supplementary Table 2). Verteporfin treatment was included as a positive control, given its previously established role in inhibiting gliomasphere migration and TEAD-dependent transcription ^16^. Thus, pharmacological inhibition of TEAD palmitoylation with VT103 or VT104 impairs the ability of GBM cells to migrate early on, and then later also impairs their ability to proliferate. These functional tumor deficits correspond to suppression of TEAD-target gene expression across different patient-derived GBM cell lines.

**Figure 1.**
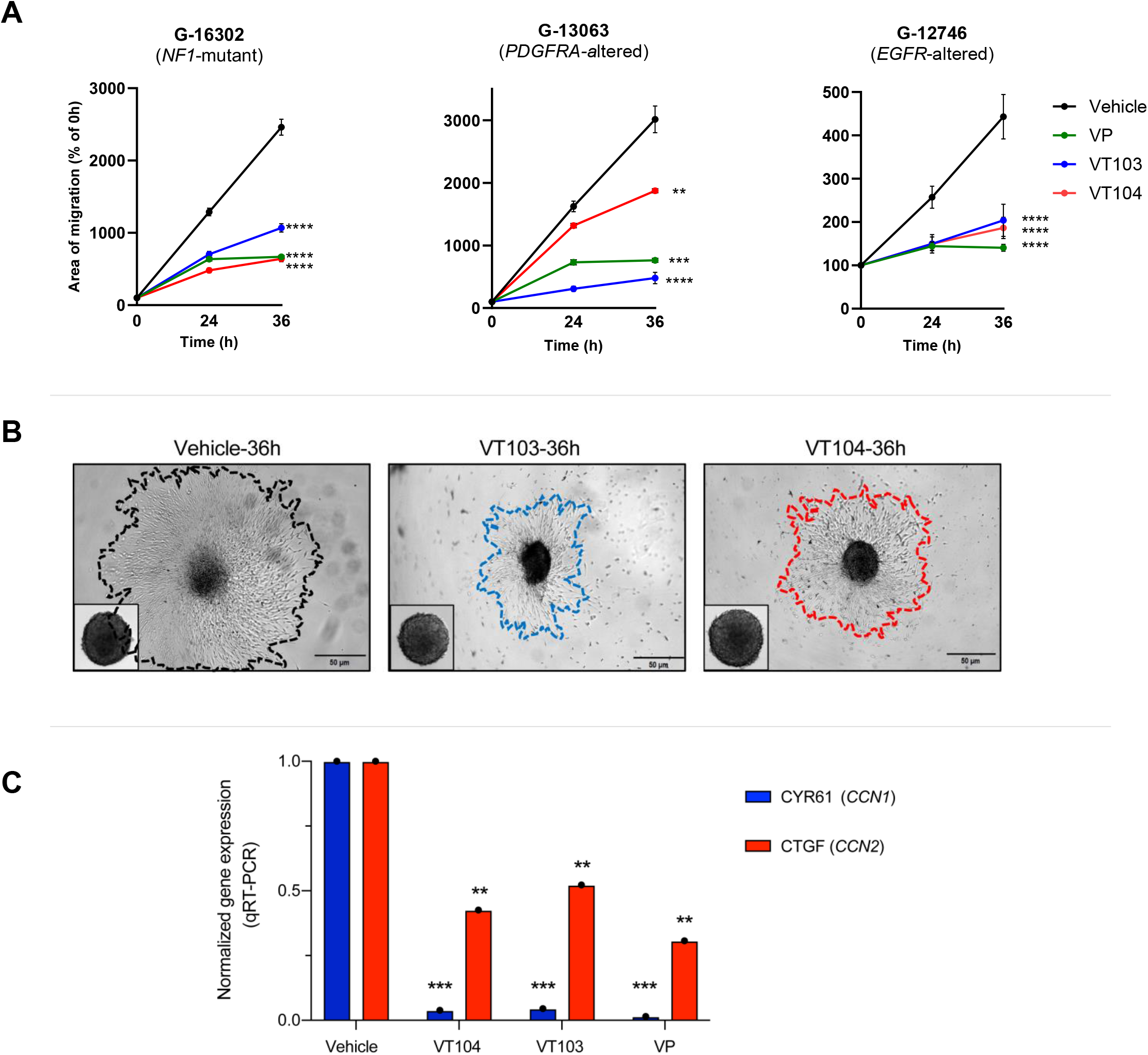
VT103 and VT104 treatment decreases GBM migration and TEAD-target expression across diverse patient-derived GBM cell lines **A.** Spheroid migration analysis shows significant decrease in confluent area of migration across three genomically distinct patient-derived GBM cell lines (*NF1*-mutant G-16302, *PDGFRA*- amplified G-13063, *EGFR*-altered G-12746) after treatment with VT103, VT104, and Verteporfin (positive control) compared to vehicle control, at 24h and 36h after drug treatment (IC_50_ dose). Each data point represents average area of spheroid migration (n=6 technical replicates), normalized to 0h time point. See also Supplementary Figures S1. **B.** Representative photomicrographs of G-13063 spheroid migration area at 0h (inset) and 36h post-treatment. **C.** VT103/VT104 pharmacodynamic analysis for TEAD-dependent transcriptional activity by qRT- PCR in G-13063 gliomaspheres from (A), 36h post-treatment. A significant decrease in TEAD- target gene expression, *CYR61* (*CCN1*) and *CTGF* (*CCN2*), is seen following treatment with VT104 and VT103 TEAD palmitoylation inhibitors, as well as with YAP-TEAD inhibitor Verteporfin used as positive control, compared to vehicle. Bars represent average gene expression normalized to *18S* housekeeping gene and to vehicle treatment (2^-ΔΔCt^) from n=3 technical replicates. **A, C**: Data are presented as mean ± SEM; statistical significance determined by two-way ANOVA (**p<0.01, ***p<0.001, ****p<0.0001).

### VT agents display robust and sustained brain penetrance

To establish the translational utility of TEAD palmitoylation inhibitors in the CNS, we next characterized the pharmacokinetic profiles of VT103 and VT104 in G-16302 PDX mice with high- grade infiltrative gliomas, after seven weeks of oral drug administration. In order to estimate true parenchymal drug penetrance, brain tissue was perfused to eliminate stagnant blood within the cerebral vasculature, immediately after plasma collection and before tissue harvesting (Figure 2A). Mass spectrometry analysis of plasma drug levels confirmed robust oral bioavailability of both VT agents acutely, 4 hours after the last drug administration, with 1362 ng/mL and 1108 ng/mL average plasma concentration for VT103 and VT104, respectively (Figure 2B, Supplementary Figure 2A). At 24 hours, plasma bioavailability for VT104 was sustained to 97% of acute levels (1077 ng/mL average plasma concentration) whereas VT103 plasma concentration decreased to 39% (Figure 2B). Notably, mass spectrometry analysis of perfused brain tissues revealed greater concentration of both VT103 and VT104 in brain compared to plasma. Specifically, acute brain:plasma percent ratios of drug (at 4h after last administration) reached 440% for VT103 and 396% for VT104. By 24 hours after last administration, brain concentration levels for both VT103 and VT104 plateaued to ∼2000ng/g, with brain:plasma ratios at or exceeding 150% (Figure 2B), indicating exceptional blood-brain barrier permeability and sustained CNS bioavailability. Further PK analysis revealed significantly longer elimination half- life (*t* _½_) of VT104 in brain tissue (∼ 70.8h) compared to VT103 (∼13.2h) (Supplementary Figure 2B), indicative of greater drug stability and slower parenchymal clearance. Importantly, systemic toxicity was not detected with either drug (Supplementary Figure 3), despite the drugs’ high brain concentration and overall long half-life kinetics. Given the stronger pan-TEAD potency of VT104 ^25^ and its more stable CNS kinetics profile compared to VT103, we prioritized VT104 in subsequent efficacy and survival studies.

**Figure 2.**
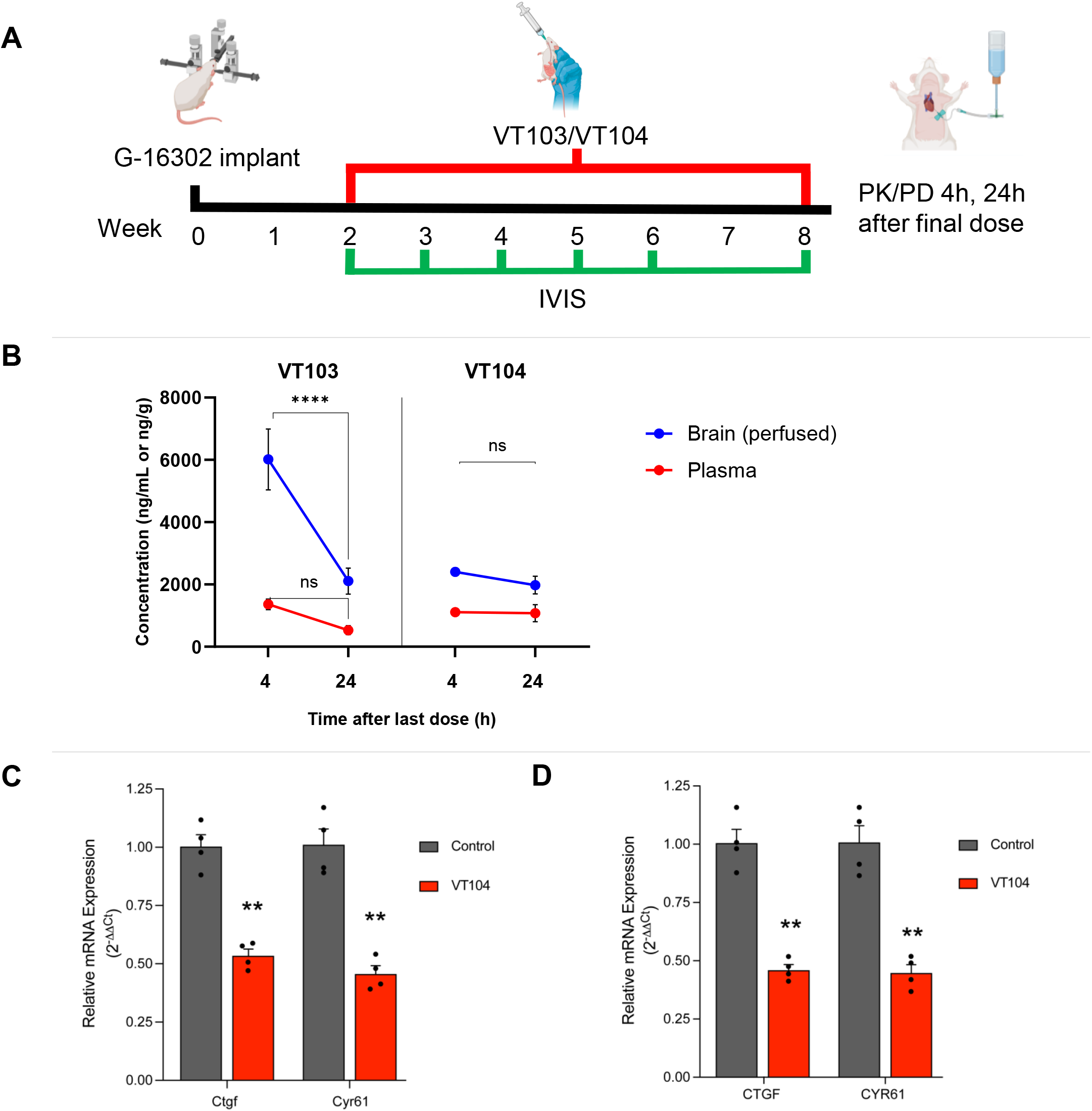
Sustained brain penetrance and TEAD inhibitor activity in-vivo of VT103 and VT104 **A.** Experimental design for VT103/VT104 pharmacokinetic studies in G-16302 PDX mice. Drug concentration is measured using mass spectroscopy 4h and 24h after last drug dose in plasma and in perfused brain tissue (n=4 mice per group). See also Supplementary Figures S2 and S3. **B.** Concentration of both VT103 and VT104 is higher in perfused brain tissue (blue) compared to plasma (red) at 4h and 24h following final dose administration, with sustained brain:plasma ratio of 396% (VT103) and 184% VT104 at 24h. VT104 maintains stable concentration in brain and plasma without statistically significant decline between timepoints (ns). VT103 levels undergo faster clearance between 4 and 24h (****, p<0.0001). Data points represent average drug concentration, and error bars represent SEM (n=4 mice, per group). Statistical significance determined by two-way ANOVA. **C.** Analysis of TEAD-target gene expression (*Ctgf*, *Cyr61*) in mouse liver tissue harvested from G- 16302 PDX mice after 7 weeks of drug treatment. **D.** Analysis of TEAD-target gene expression (*CTGF*, *CYR61*) in xenograft human gliomas harvested from G-16302 PDX brains after 7 weeks of drug treatment. **C, D**. Bar plot represents averaged gene expression normalized to *Rpl13a* (C) or *RPLPO* (D) housekeeping genes and to control vehicle treatment, using species-specific primers (n=4 mice). Data are presented as mean ± SEM; statistical significance determined by two-tailed Student’s t-test (**p<0.01).

### VT104 monotherapy delays GBM progression in-vivo

Upon establishing broad anti-GBM properties in-vitro and robust CNS penetrance in-vivo, we focused the remainder of this study on the translational potential of VT104 for patients with recurrent GBM, a demographic that lacks effective treatment and would be eligible for repurposed drug therapy in prospective clinical trials. First, we confirmed pharmacodynamic (PD) properties of VT104 in-vivo, in the same G-16302 PDX mice used for PK studies. We used species- specific qRT-PCR primers (Supplementary Table 2) to measure the effect of treatment on TEAD- mediated gene expression and detected significant downregulation of TEAD-target genes *CTGF* and *CYR61* both in control mouse liver (Figure 2C) and in human PDX brain tumors (Figure 2D) after 7 weeks of VT104 administration, compared to vehicle-treated controls. This confirmed that the high concertation of VT104 within the brain results in on-target inhibition of YAP-TEAD activity. Next, we tested the efficacy of VT104 as monotherapy, choosing the orally administered dose 10mg/kg, as this dose displayed excellent PK/PD profile in the CNS and has been previously shown to be therapeutic without causing significant toxicity in non-glioma mouse models ^25–27^. To monitor the impact on tumor burden, we utilized the recurrent GBM orthotopic PDX model, G-16302_akaluc, expressing the Akaluc bioluminescent reporter. We have previously shown superior bioluminescence of Akaluc over standard luciferase reporter in orthotopic GBM PDX models and have successfully used G-16302_akaluc for non-invasive monitoring of GBM tumor burden over time ^32^. Two weeks after G-16302_akaluc tumor implantation, PDX mice were treated with oral VT104 or vehicle for an additional seven weeks. Serial bioluminescence imaging (BLI) demonstrated that while all tumors exhibited progressive growth, continuous VT104 treatment resulted in delayed tumor progression, where VT104-treated animals showed significantly lower bioluminescent tumor flux compared to vehicle control mice at week 8 (Figure 3A-B). The results suggest a therapeutic window during which pharmacological disruption of YAP- TEAD activity constrains rapid tumor growth, effectively delaying the characteristic tumor progression of recurrent GBM.

**Figure 3.**
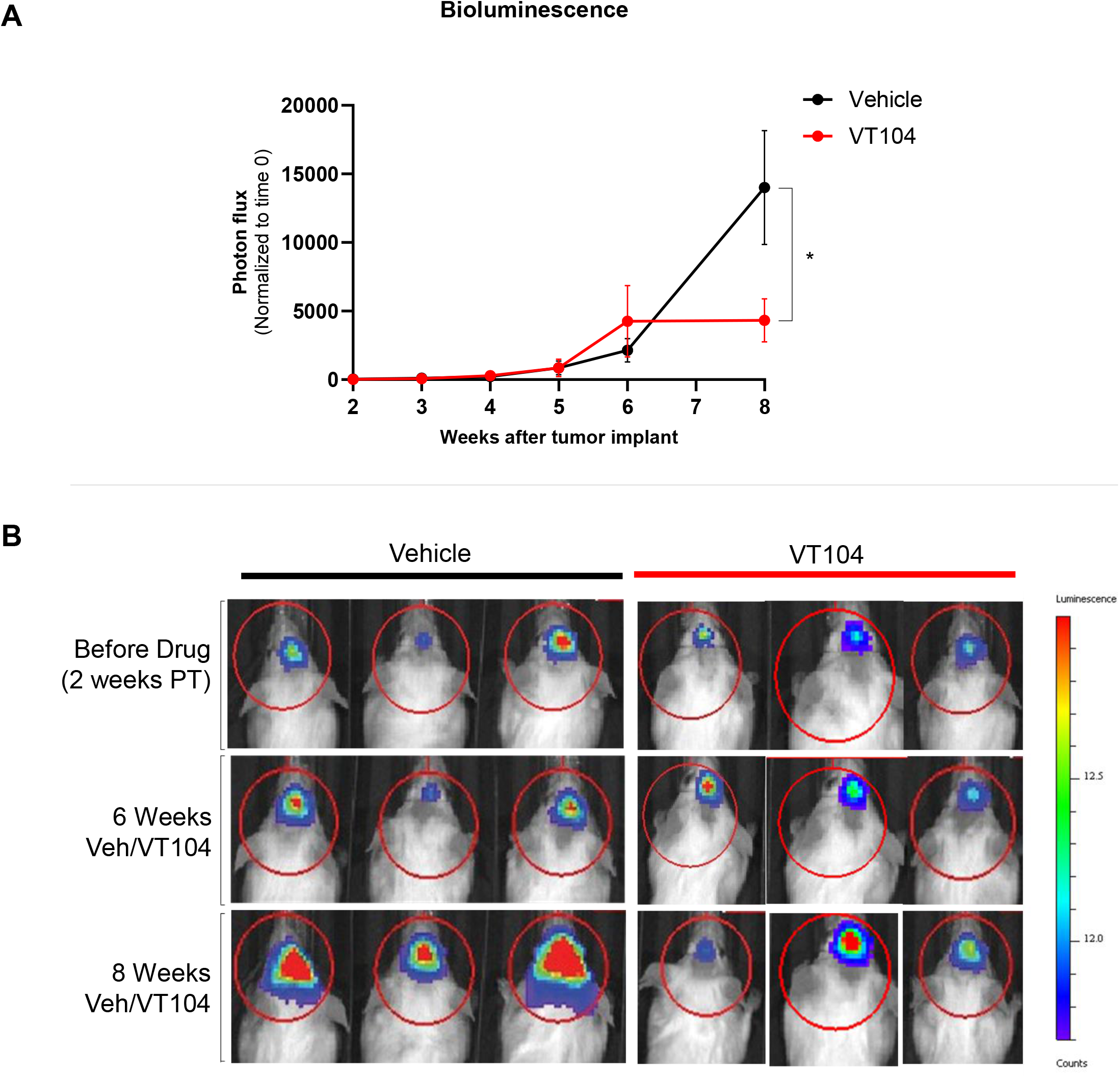
Oral VT104 monotherapy delays GBM progression in-vivo **A.** Longitudinal bioluminescence imaging (BLI) study to assess tumor growth in anaesthetized G- 16302_akaluc mice with orthotopic PDX tumors, treated with either vehicle (control) or oral VT104 (10mg/kg) continuously for 7 weeks, beginning 14 days after tumor transplantation. A significantly lower photon flux from tumors is seen in VT104-treated mice compared to control on week 8 (*p=0.029) (n=8 mice per group for weeks 2-6; n=6 mice per group for week 8). Data points represent mean photon flux (photons/sec) from Akaluc bioluminescence, normalized to 0-min signal; error bars represent SEM. Statistical significance determined by two-tailed Student’s t-test. **B.** Representative images of progressive BLI signal in tumors of anaesthetized G-16302_akaluc mice treated with either vehicle or VT104, from part A. PT = post-transplantation of tumor.

### VT104 therapy combined with chemoradiation extends survival in recurrent GBM models

Given the observed therapeutic window of VT104, we conducted long-term survival studies to assess the efficacy of VT104 in preclinical GBM models, as either monotherapy or in combination with radiotherapy and temozolomide (RT+TMZ) (Figure 4A). Studies were performed using two genetically distinct recurrent GBM PDX models: G-16302 (*NF1*-mutant, *EGFR*-wildtype) and G- 12315 (*EGFR*-amplified/mutant), generated from patients who recurred after chemoradiation ^31^ and in which GBM cells are resistant to RT+TMZ ^16^. In the *NF1*-mutant G-16302 model, combination VT104+RT+TMZ treatment resulted in significantly prolonged survival compared to vehicle control, to VT104 monotherapy, or to RT+TMZ treatment groups (Figure 4B; *p=0.0291, log-rank test). Treatment with RT+TMZ alone failed to show significant survival difference compared to control, consistent with the expected biology of this cell line ^16, 31^. Despite showing a transient delay in tumor progression, treatment with VT104 alone did not prolong survival in this aggressive, recurrent GBM model. The therapeutic efficacy with combined VT104+RT+TMZ was reproduced in a second, genetically distinct *EGFR*-amplified G-12315 recurrent GBM model, in which the conferred survival benefit was more pronounced (Figure 4C). In this highly aggressive background, the addition of VT104 to RT+TMZ conferred a highly significant survival benefit compared to other treatment groups (***p = 0.0001, log-rank test). Importantly, tumor implantation was confirmed in PDX mice, and no signs of systemic toxicity were seen after prolonged, continuous treatment with VT104. Together, these data suggest that VT104 is a highly effective CNS therapeutic when combined with standard of care RT+TMZ, significantly extending survival in two molecularly distinct subtypes of recurrent GBM.

**Figure 4.**
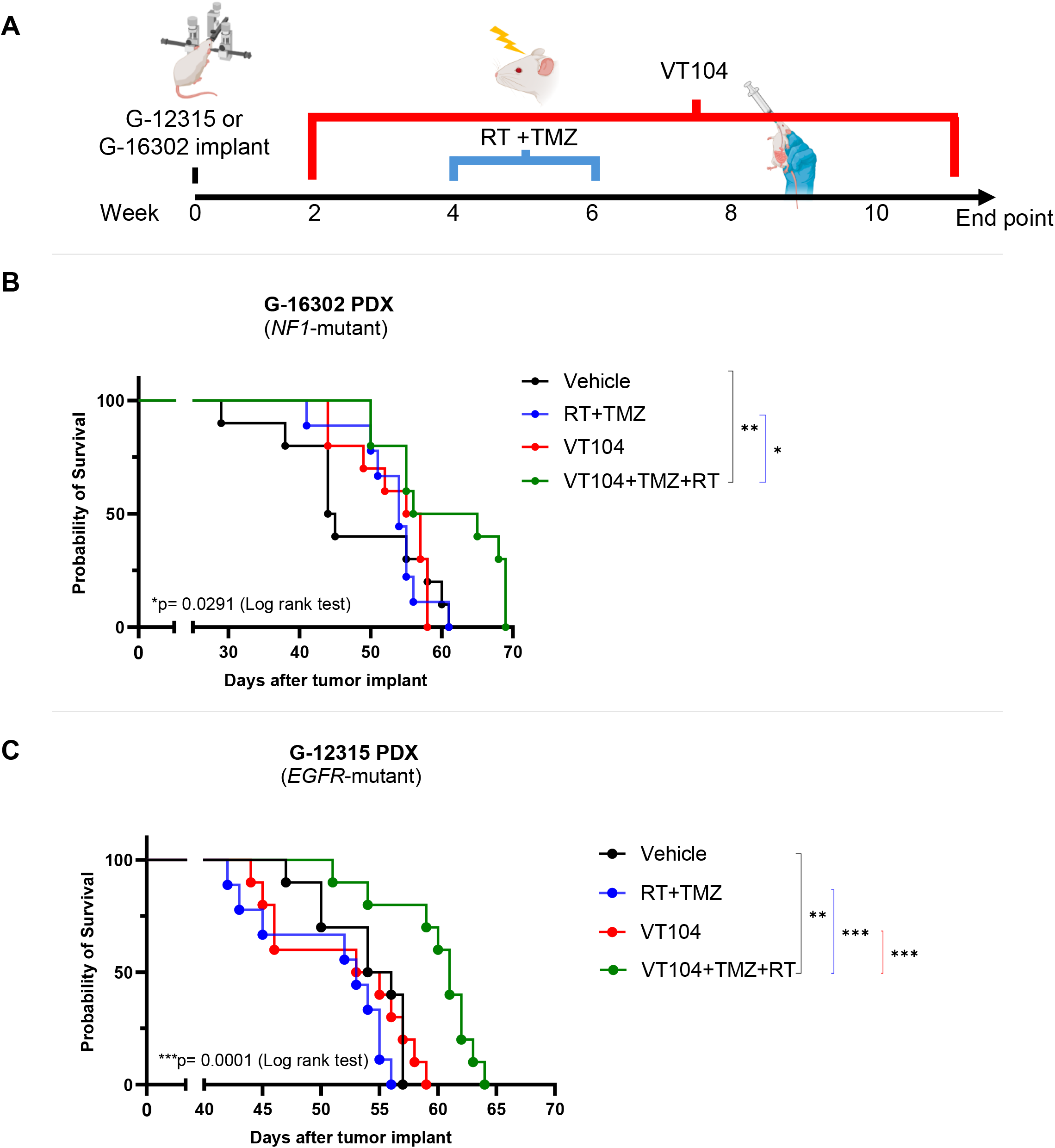
Oral VT104 confers survival efficacy in combination with chemoradiation in two recurrent GBM models **A.** Schematic of treatment regimen for mice with G-16302 or G-12315 recurrent GBM PDX. **B.** Kaplan-Meier survival analysis of *NF1*-mutant G-16302 PDX recurrent GBM model (n=10 mice per group). Combined VT104+RT+TMZ therapy significantly extends overall survival compared to other cohorts (*p=0.029, by global log-rank test; **p=0.0085 vs. vehicle control, *p=0.014 vs. RT+TMZ, by pairwise comparisons). **C.** Kaplan-Meier survival analysis of *EGFR*-amplified G-12315 PDX recurrent GBM model (n=10 mice per group). Combined VT104+RT+TMZ therapy significantly extends overall survival compared to other cohorts (***, p=0.0001, by global log-rank test; **p=0.0012 vs. vehicle control, ***p=0.0006 vs. VT104, ***p=0.0005 vs. RT+TMZ, by pairwise comparisons).

### VT104 therapy combined with chemoradiation diminishes infiltrative tumor burden in *EGFR*- mutant recurrent GBM

To evaluate the impact of the different therapeutic regimens on overall tumor burden and on extent of GBM infiltration, we performed histopathological analysis in the G-12315 PDX model, where survival benefit from combined VT104+RT+TMZ treatment was most significant. This confirmed significant decrease in overall and infiltrative tumor burden (Figure 5A-E), concordant with the observed survival efficacy. Vehicle-treated control animals (Figure 5A) exhibited expansive, hypercellular tumor cores with associated ventricular compression, consistent with mass effect on the adjacent brain parenchyma. Similar patterns of aggressive growth and extensive tumor infiltration were observed in the VT104 alone (Figure 5B) and the RT+TMZ (Figure 5C) treatment groups. In contrast, triple combination therapy (VT104+RT+TMZ) demonstrated a noticeable and statistically significant reduction in tumor burden (Figure 5D-E). Some G-12315 PDX mice receiving combination therapy displayed minimal residual neoplastic infiltration, with a notable preservation of the surrounding neuroanatomical architecture and a marked absence of prominent mass-like tumor cores observed in other experimental groups (Figure 5D). Overall, these findings suggest that VT104 can act synergistically with RT+TMZ to improve survival in *EGFR*-mutant recurrent GBM, potentially by sensitizing standard-of-care- resistant, recurrent GBM cells to radiotherapy and/or temozolomide.

**Figure 5.**
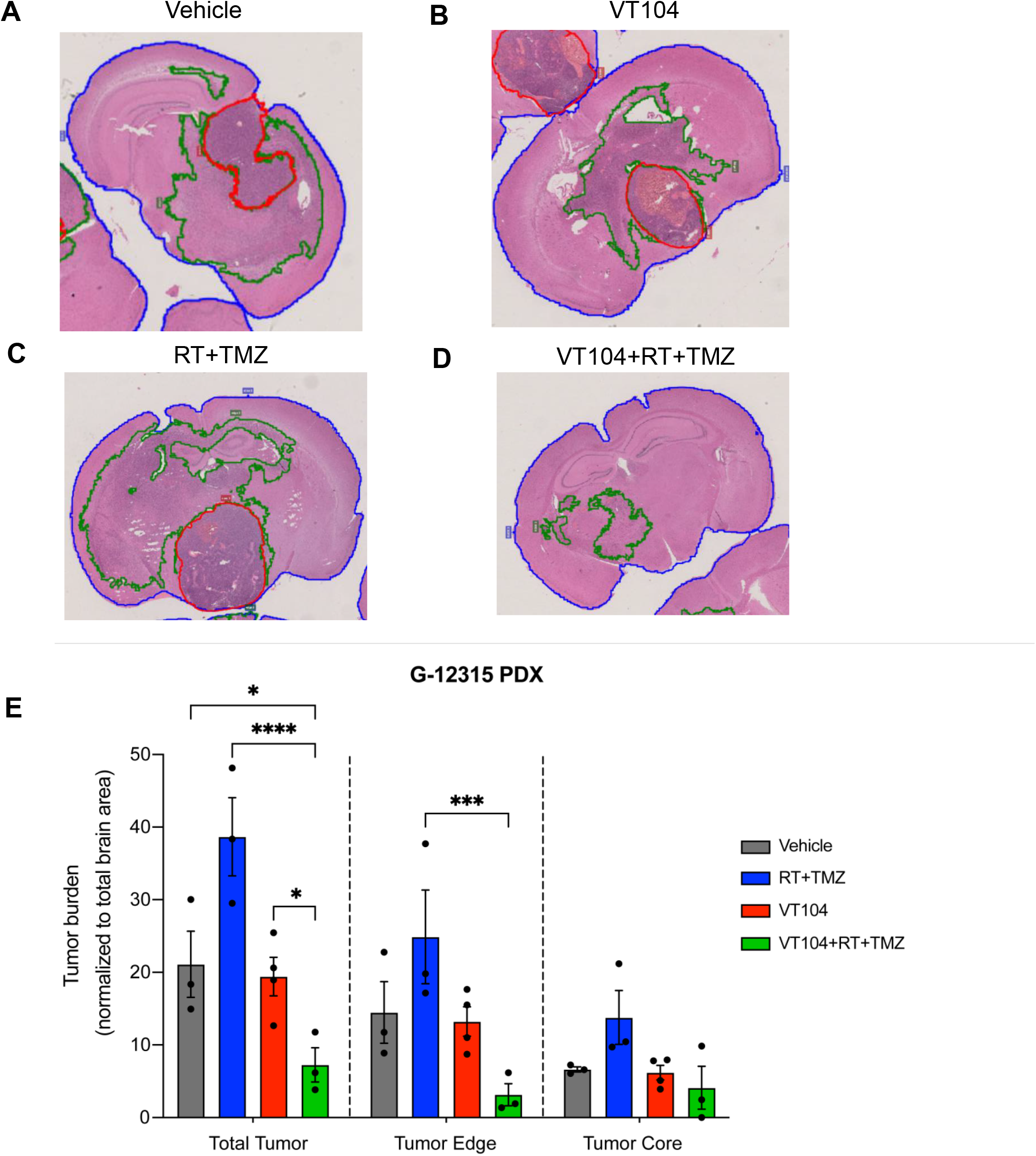
VT104 therapy combined with chemoradiation decreases infiltrative tumor burden in *EGFR*-mutant PDX mice **A-D.** Representative photomicrographs of H&E-stained coronal brain sections of G-12315 PDX mice, obtained at survival endpoint for each experimental group (A: vehicle; B: VT104; C: RT+TMZ; D: VT104+RT+TMZ). Coronal sections with largest representative area of tumor burden are evaluated for each mouse, quantifying area of tumor core (solid red line with highest (>80%) tumor cell density, near tumor implantation site, showing tumor necrosis and/or vascular proliferation) and infiltrative tumor edge (solid green line, lower tumor density with diffuse infiltration, lacking tumor necrosis or vascular proliferation). **E.** Bar graph quantification of tumor burden in G-12315 survival cohort mice across experimental groups, showing significantly lower overall (total) and infiltrative edge tumor area in mice receiving combined VT104+RT+TMZ therapy (n=3-4 mice per group, n=2-3 coronal sections per mouse). For each coronal section, tumor area is normalized to total brain area. Data are presented as mean ± SEM; statistical significance determined by two-way ANOVA with only significant differences shown (* p<0.05, *** p=0.0004, **** p<0.0001).

### VT104 therapy combined with chemoradiation potentiates genotoxic stress and TEAD inhibition in GBM cells

To investigate potential mechanisms of synergistic action between VT104 and chemoradiation, we performed reductionistic functional in-vitro studies, leveraging the stronger responder G- 12315 recurrent GBM cells (Figure 6A). To tease out effects from each treatment group, we included treatment with VT104, TMZ, or irradiation (IR) alone, as well as all double and triple treatment combinations. We then assessed the effects of therapy on genotoxic stress and cytotoxicity by measuring DNA double strand damage, a known effect of IR and of alkylating TMZ chemotherapy ^42^, as well as VT104’s on-target effects on downstream TEAD activity, a known molecular driver of GBM cell migration ^17^. In addition to the canonical TEAD-target genes, *CTGF* and *CYR61,* we also included *EGFR* as a TEAD1-regulated, pro-migration marker ^17^, part of a pan- GBM tumor periphery infiltration gene signature ^34^.

**Figure 6.**
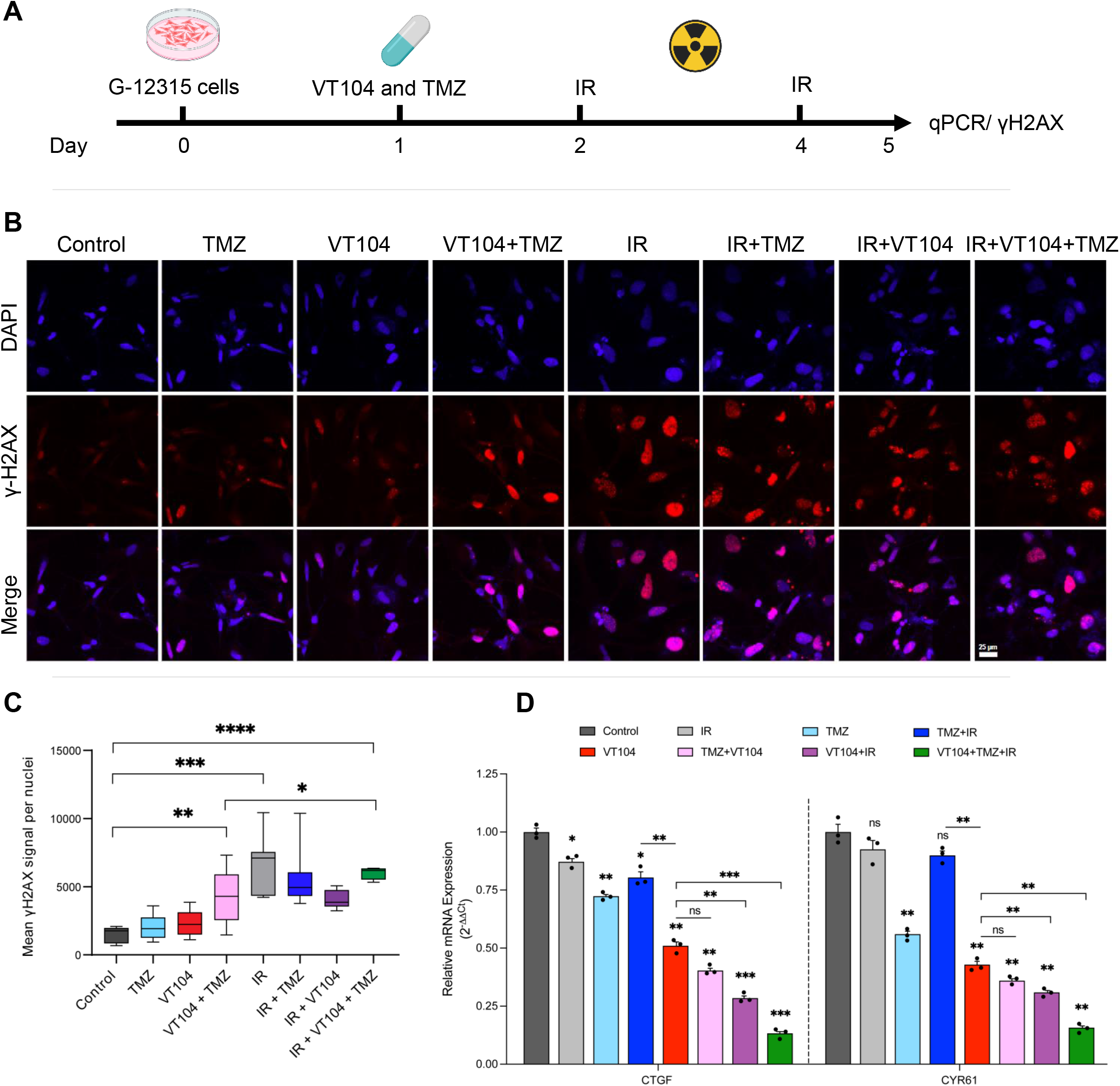
VT104 therapy combined with chemoradiation potentiates genotoxic stress and TEAD suppression in recurrent GBM cell lines **A.** Schematic of experimental design. G-12315 cells are treated with VT104 (20 μM), TMZ (50 μM) and/or IR (2Gy+2Gy) and harvested 96 hours later for immunofluorescence and qPCR analyses. **B.** Representative immunofluorescence images of γ-H2AX intensity as a marker of double stranded DNA damage and DAPI used as nuclear stain. **C.** Box and whiskers plot showing quantification of γ-H2AX intensity per nucleus (n ≥50 nuclei in ≥4 fields). Line marks the median and whiskers mark the maximum and minimum values. Statistical significance determined by two-tailed t-test with Welch’s correction (**** p<0.0001, *** p<0.001, ** p<0.01, * p<0.05). **D.** Analysis of TEAD-target gene expression (*CTGF*, *CYR61*) in G-12315 PDX cells shows strongest downregulation after triple combination (VT104+TMZ+RT). Bar plot represents averaged gene expression normalized to housekeeping *18S* gene. Data are presented as mean ± SEM (n=3 wells per treatment group). Statistical significance determined by two-way Student’s t-test (*p < 0.05, **p < 0.01, ***p < 0.001). See also Supplementary Figure S4A.

To assess cellular effects on genotoxic stress, we measured intensity of nuclear γ-H2AX, a known marker of double stranded DNA damage (Figure 6B-C). We saw expected increase in γ-H2AX intensity in cells treated with IR, compared to non-IR treated. Of note, TMZ treatment by itself did not show a significant increase γ-H2AX levels compared to control (Figure 6C), consistent with the presumed chemoresistance of G-12315 GBM cells derived after failed TMZ treatment and tumor recurrence. However, combination of VT104+TMZ displayed significantly higher γ-H2AX intensity in cells compared to TMZ alone, equivalent to levels seen after IR treatment. Triple combination therapy (VT104+TMZ+IR) displayed significantly higher γ-H2AX intensity compared to VT104+TMZ. These cells demonstrated morphological signs of cytotoxicity including nuclear fragmentation with micronuclei and nuclear shrinkage with chromatin condensation and apoptotic blebs (Figure 6B). This analysis suggests the potential role of VT104 in re-sensitizing TMZ-resistant cells to DNA double stranded break and genotoxicity.

Evaluation of TEAD activity in these groups afforded additional rationale for the observed synergy. First, treatment of recurrent GBM G-12315 cells with VT104 alone demonstrated expected downregulation of TEAD activity (Figure 6D, red bar), confirming previous PD data seen with other patient-derived GBM cell lines. Double treatment of GBM cells with VT104+TMZ (pink bar) or with VT104+IR (purple bar) downregulated TEAD-target gene expression to a similar or slightly greater extent compared to VT104 treatment alone. Notably, triple combination treatment with VT104+TMZ+IR demonstrated the greatest inhibition of TEAD activity, with robust downregulation of *CTGF* and *CYR61* expression to nearly 10% of control (Figure 6D, green bar) and downregulation of *EGFR* expression to approximately 20% of control (Supplementary Figure 4A, green bar). While IR treatment (light gray bar) did not affect TEAD activity to a significant extent, treatment of cells with TMZ (light blue) and to a smaller extent TMZ+IR (dark blue bar) did show mild off-target effect on TEAD activity as well, which was significant compared to control but also significantly less compared to VT104 treatment alone (Figure 6D).

The observed synergistic effects on TEAD activity and *EGFR* expression were intriguing and motivated additional G-12315 PDX studies to confirm if similar effects are seen within the constraints of physiological drug penetrance in-vivo. We recapitulated all treatment groups, and treated animals with a similar drug dose as previously used in survival studies; harvested tissue was prioritized for TEAD-target gene molecular analysis rather than end-point survival or histology (Figure 7A). With this analysis, we first confirmed VT104 pharmacodynamics in our G- 12315 PDX in-vivo models, detecting approximately 50% reduction of TEAD activity after treatment alone (Figure 7B, red bar). Notably, VT104+TMZ+RT combination therapy showed a profound inhibitory effect on TEAD activity, corroborating prior in-vitro findings, with more than 90% downregulation of *CTGF* and *CYR61* (Figure 7B, green bars) and strong downregulation of *EGFR* (Supplementary Figure 4B), all of them being significantly lower than VT104, VT104+RT, and VT104+TMZ treatments. In contrast, double combination therapy, VT104+RT and VT104+TMZ, had an equivalent effect on TEAD activity compared to VT104 therapy alone (Figure 7B, pink/purple vs. red bars).

**Figure 7.**
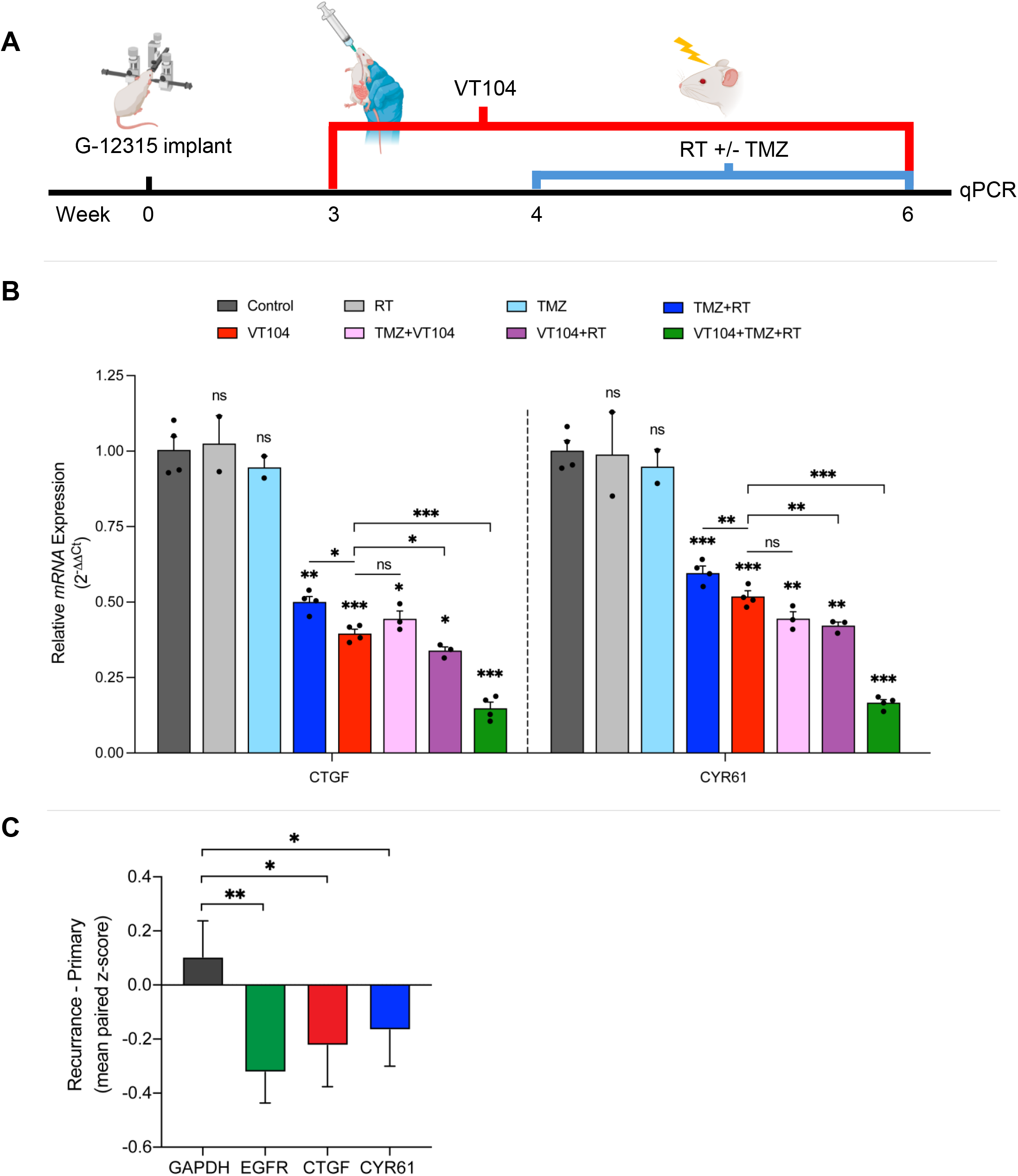
Chemoradiation combined with VT104 therapy augments TEAD suppression in recurrent GBM **A.** Schematic of experimental design and treatment groups. **B.** Analysis of TEAD-target gene expression (*CTGF*, *CYR61*) in human gliomas harvested from G- 12315 PDX mice, showing greatest downregulation after triple combination therapy (VT104+TMZ+RT). Treatment began three weeks after tumor implantation (VT104: 10mg/kg/day p.o. for 3 weeks; TMZ: 25mg/kg/day p.o. for 5+5 days; RT: 2Gy/day for 5+5 days, 10Gy total). Bar plot represents averaged gene expression normalized to housekeeping *RPLPO* gene. Data are presented as mean ± SEM (n=4 mice per treatment group). Statistical significance determined by two-way Student’s t-test (*p < 0.05, **p < 0.01, ***p < 0.001). See also Supplementary Figure S4B. **C.** Bar graph representations of TEAD-target gene RNA-seq expression analyses in paired primary-recurrent IDH-wildtype GBM patient samples obtained from the longitudinal GLASS consortium (n=68 primary tumor samples with no treatment, n=68 patient-matched first recurrences, RT+TMZ treated). Data are presented as mean ± SEM. Statistical significance determined by one-way Student’s t-test (**p=0.003 for *EGFR*; *p=0.03 for *CTGF* and *CYR61*). See also Supplementary Figure S5.

Unexpectedly, we detected a significant downregulation of TEAD activity after TMZ+RT chemotherapy, albeit less pronounced compared to treatment groups receiving VT104 (Figure 7B, blue vs. red bars). To investigate further the clinical relevance of this non-specific (off-target) TEAD inhibitory effect by standard chemoradiation, we analyzed the GLASS consortium’s large RNA-seq dataset of paired primary-recurrent IDH-wildtype GBM patient samples ^35, 36^, selecting primary GBM resections without treatment and patient-matched first recurrences after treatment with both TMZ+RT. Analysis confirmed significant downregulation of *CTGF*, *CYR61*, and *EGFR* after TMZ+RT, corroborating in-vivo PDX results (Figure 7C). Several other TEAD-target genes were also noted to be preferentially downregulated after TMZ+RT treatment, which was independent of EGFR-amplification status (Supplementary Figure 5).

Overall, this analysis puts forward both genotoxic stress and suppression of oncogenic TEAD activity as two biological phenomena underlying the observed synergistic efficacy of VT104 when combined with chemoradiation, with genotoxic stress having primarily a TMZ-related sensitization effect while TEAD inhibition being augmented by both TMZ and RT. The results suggest that chemoradiation greatly potentiates the dynamics of VT104 as an inhibitor of TEAD- mediated transcription, and vice versa, a novel finding with important implications for current and future combination therapies with VT inhibitors.

## Discussion

The management of recurrent GBM remains one of the greatest challenges in neuro-oncology. Despite aggressive standard-of-care regimens ^5^, the disease is characterized by a persistent therapeutic plateau, driven by diffuse parenchymal infiltration and the rapid emergence of adaptative resistance ^43, 44^. YAP-TEAD signaling has been recently shown to be a central, convergent mechanism of maintaining adaptive GBM stem-like plasticity and therapeutic resistance ^11^. In the present study, we investigated the role of TEAD palmitoylation inhibitors as rational targeted inhibitors of therapeutic resistance in preclinical models of recurrent GBM. Our study establishes that VT103 and VT104 TEAD inhibitors are extremely brain penetrant, a novel and translationally relevant finding in the context of future clinical studies. Moreover, we discover that VT104, a potent and orally bioavailable inhibitor of pan-TEAD palmitoylation ^25^, acts synergistically with RT and TMZ to improve overall survival in standard-of-care-resistant recurrent GBM PDX models, and that this synergism is at least partially mediated through the potentiation of genotoxic stress and through the inhibition of oncogenic TEAD activity. Targeting the YAP-TEAD axis with VT104 not only suppressed tumor growth, but it also impeded tumor infiltration, underscoring adjuvant VT drug therapy as a clinically viable therapeutic strategy for future GBM treatment efforts.

### Exceptional CNS penetrance

A major limitation of targeted therapies in GBM is the inadequate penetration of drugs across the blood-brain barrier, resulting in subtherapeutic CNS exposure. Prior Hippo pathway inhibitors, such as Verteporfin ^12^, demonstrated preclinical efficacy in reducing tumor proliferation and invasion ^15, 16, 19–21^ but have been limited for clinical translation due to difficulty in outpatient administration, poor solubility, and suboptimal CNS penetration ^22^. In contrast, VT103 and VT104 were rationally designed as small molecules of approximately 420 Da in size and lipophilic in nature ^25^, thus having potentially superior blood-brain barrier penetrance through passive diffusion. Indeed, our PK analysis demonstrates that VT103 and VT104 achieve brain:plasma (B:P) ratios exceeding 150% even at 24h after administration, indicating robust CNS exposure. Importantly, drug concentration in brain tissue was measured after perfusion, thus eliminating any potential plasma drug contamination. Moreover, this sustained brain bioavailability was seen after oral administration, underscoring the superior translational potential for these TEAD inhibitors. Higher drug concentration in brain compared to plasma (B:P ratio greater than 100%) is rarely encountered with brain chemotherapeutics and is considered exceptionally brain penetrant ^45^. The B:P ratio of Temozolomide is only ∼ 18% (0.18) ^46^ and it is considered acceptable and standard-of care. VP’s highest reported ratio is 50% (0.5) ^22^ and the antidepressant mirtazapine has a B:P ratio of 153% (1.53) ^47^. Thus, VT drugs have much higher brain biodistribution than TMZ and VP, similar to that of antidepressants, making them remarkable CNS candidate chemotherapeutics from a chemical perspective. Importantly, VT104 inhibited TEAD-target gene expression in both of our in-vivo orthotopic PDX models, confirming that the drug was not only bioavailable in the brain but that it was pharmacologically active. The favorable PK/PD properties of VT agents, combined with the survival efficacy seen for VT104, confer important translational advantage and support the feasibility of inhibiting TEAD palmitoylation as a therapeutic strategy for recurrent GBM.

### TEAD palmitoylation is a targetable vulnerability in EGFR-driven recurrent GBM

To address the biological heterogeneity of GBM, we evaluated VT104 efficacy in molecularly distinct PDX models. Notably, VT104 displayed pronounced anti-invasive GBM response and conferred greatest survival efficacy in the *EGFR*-amplified/*EGFR*-mutant (G-12315) PDX model. This observed vulnerability of EGFR-driven GBM to YAP-TEAD inhibitors is consistent with prior studies demonstrating bidirectional crosstalk dependency between EGFR and YAP-TEAD signaling, including EGFR/RTK activation of YAP ^48–50^ and direct, TEAD-mediated regulation of *EGFR* transcription ^15–17, 34^. Nevertheless, survival benefit was also observed in the *NF1*-mutant and *EGFR* wildtype (G-16302) PDX model. Recurrent GBM tumors, and especially those that are *NF1*-mutant, are considered more mesenchymal-like, and the co-activator TAZ can act as a master regulator of mesenchymal differentiation in GBM ^51^, providing molecular rationale for the observed survival efficacy. Overall, our findings support the broad relevance of TEAD inhibitors in recurrent GBM but prioritize *EGFR*-altered recurrent GBM patients as the strongest response candidates.

### Synergy and the disruption of adaptative resistance

Despite remarkable brain penetrance, mouse efficacy studies indicated that VT104 monotherapy is insufficient to extend survival as a standalone agent in recurrent GBM and that its therapeutic value is dependent on combined administration with temozolomide and radiotherapy. These results are consistent with prior studies, showing the greatest efficacy of YAP-TEAD inhibitors when combined with radiation, chemotherapy ^11, 20, 52–55^ or immunotherapy ^19^ in GBM, and support the concept of reversing GBM’s adaptative resistance to conventional therapies through targeting the YAP-TEAD axis ^11^. Proposed mechanisms of synergy include sensitization to TMZ and radiation through apoptosis, augmented cell cycle arrest and suppression of adaptative survival pathways ^20, 11, 52^. Our study uncovered additional potential explanations for the observed synergism between TEAD inhibition and chemoradiation: the augmented suppression of oncogenic TEAD activity and the enhanced sensitivity of GBM cells to TMZ-mediated genotoxic stress when VT104 is combined with chemoradiation. Interestingly, we discovered that chemoradiation alone can suppress TEAD activity, an unexpected finding which we confirmed across a large cohort of patient samples and across several TEAD-target genes. YAP-TEAD activity has been strongly implicated in driving GBM migration and invasion ^16, 17, 56^, at least in part through the regulation of EGFR transcription ^15, 34^. Thus, the augmented suppression of TEAD activity, including *EGFR*, when chemoradiation is combined with VT104, provides a rationale for the decreased infiltrative phenotype and increased survival of *EGFR*-mutant PDX mice treated with VT104+TMZ+RT.

The observed synergistic phenomena, i.e. genotoxic stress and TEAD activity suppression, may be independent or potentially related events. In support of the latter, a recent study found that TEADs co-localize to double stranded breaks within the nuclear DNA, suggesting a potential role for TEAD in DNA damage repair ^57^. One possibility is that double-stranded DNA damage induced by RT+/-TMZ could trap TEAD, preventing its normal transcriptional activity, and effectively enhancing VT104’s inhibition of TEAD-mediated transcriptional activity. This could explain the remarkable decrease in tumor burden and survival efficacy seen in VT104+TMZ+RT mice, compared to TMZ+RT alone. The exact epigenetic mechanism of synergistic action is beyond the scope of this preclinical-focused study and deserves focused attention in future investigations.

In conclusion, VT104 therapy achieves robust CNS exposure, inhibits GBM infiltration, and enhances the therapeutic efficacy of chemoradiation treatment in recurrent GBM models by potentiating genotoxic stress and attenuating TEAD-dependent transcription related to tumor migration. These findings provide strong preclinical justification for clinical evaluation of TEAD palmitoylation inhibitors as a rational and clinically actionable therapeutic strategy for the treatment of patients with recurrent GBM.

### Limitations

While our findings demonstrate strong preclinical efficacy, several limitations must be acknowledged, First, these studies were conducted in immunocompromised mouse models, which do not fully recapitulate the immune microenvironment of human GBM. Second, although VT104 demonstrated excellent CNS penetration and survival benefit in combination with standard of care, long term safety and toxicity profiles require further investigation. Future studies in additional GBM models and clinically relevant therapeutic contexts will be essential to further establish the clinical and translational potential of VT104.

## Author Contributions

Thenzing Judá Silva Hurtado:

Overall project leadership, data acquisition and results interpretation; experimental design, data acquisition and analysis of stereotaxic intracranial injections, animal treatments, BLI imaging, survival and histology studies, cell culture; manuscript and figures generation.

Balagopal Pai, Anne Marie Barrette:

Support in experimental design, data acquisition and analysis of cell migration and viability studies; manuscript review and editing.

Mohammad Imran K. Khan:

Support in experimental design, data acquisition and analysis of qPCR and cell proliferation studies; manuscript review and editing.

Debdatta Halder:

Genotoxic stress data acquisition and analysis; manuscript review and editing.

Timothy Richardson:

GLASS consortium RNA-seq data acquisition; manuscript review and editing.

Raymund Yong:

Procurement of patient-derived tumor tissue and cell lines.

Roland H. Friedel, Hongyan Zou, Anirudh Sattiraju:

Generation and technical expertise with G-16302_Akaluc cell line for BLI experiments

Dolores Hambardzumyan:

Expertise with mouse radiotherapy and tumor microenvironment.

Tracy Tang:

Supply of VT compounds; input on experimental design and PK analysis.

Nadejda M. Tsankova:

Conception of project and primary supervision; experimental design oversight; final manuscript generation, review and editing.

## Consent for publication

All authors have approved the manuscript for publication.

## Statement of translational relevance

Resistance to therapy causes rapid progression and clinical decline in recurrent GBM patients, highlighting the urgent need for novel strategies to target adaptative plasticity. By integrating YAP-TEAD inhibition with standard-of-care therapy, we provide a clinically relevant paradigm for synergistic treatment in recurrent GBM that both inhibits tumor infiltration and effectively reduces therapeutic resistance to chemoradiation. We demonstrate that the TEAD palmitoylation inhibitor VT104 robustly penetrates the blood brain barrier and that it has a significant survival benefit in combination with radiotherapy plus temozolomide in two preclinical recurrent GBM models. VT compounds have already demonstrated favorable tolerability in the ongoing NCT04665206 Clinical Trial for mesothelioma and solid metastases, and our study results motivates their use as adjuvant therapy in recurrent GBM patients. Importantly, VT TEAD inhibitors are small molecule drugs that can be administered as oral pills in an outpatient setting, improving patient compliance, quality of life, and health care cost.

## Acknowledgements

We thank the Center for Comparative Medicine and Surgery (CCMS) at the Icahn School of Medicine at Mount Sinai for their assistance with animal care and housing.

## SUPPLEMENTARY DATA

**Supplementary Table 1.**
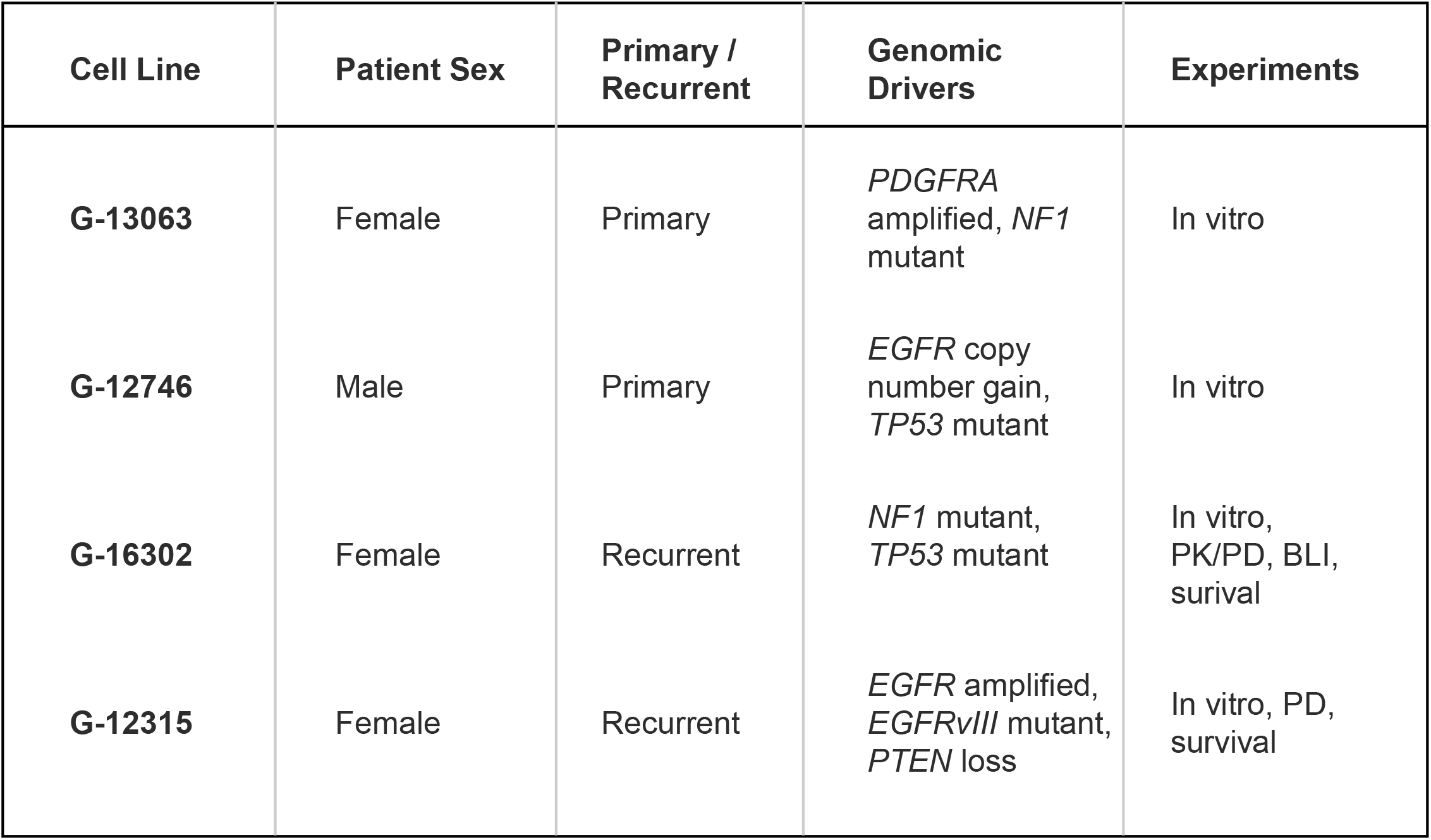
(Relates to all Figures) Clinical and molecular characteristics of patient-derived glioblastoma cell lines Information summary for all patient-derived IDH-wildtype GBM cell lines used, including patient sex, tumor genomic drivers, primary/recurrent tumor status, and experiment use.

**Supplementary Table 2.**
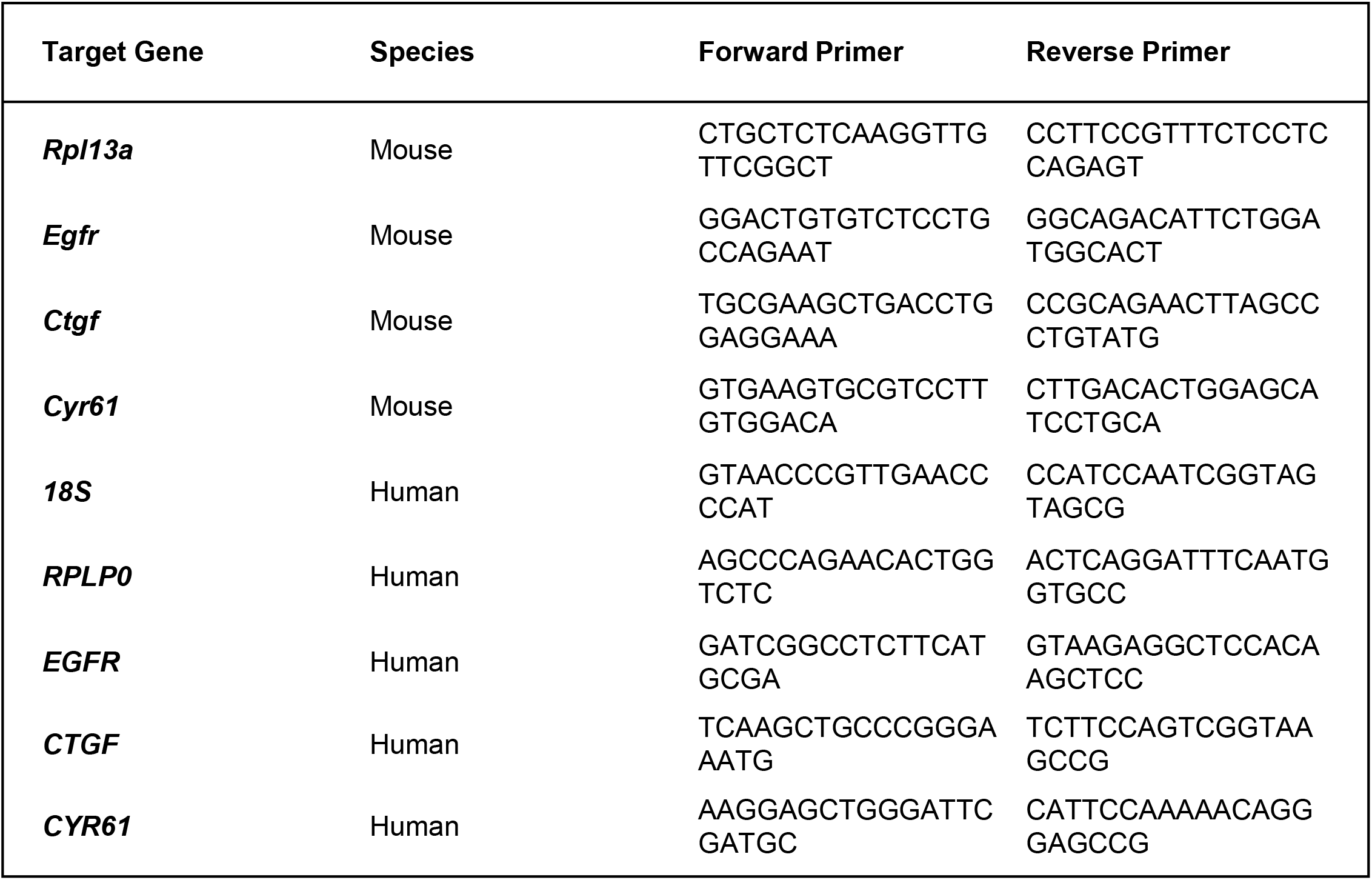
(Relates to Figures 1, 2, 6, 7) Summary of species-specific qRT-PCR primers

**Supplementary Figure S1.**
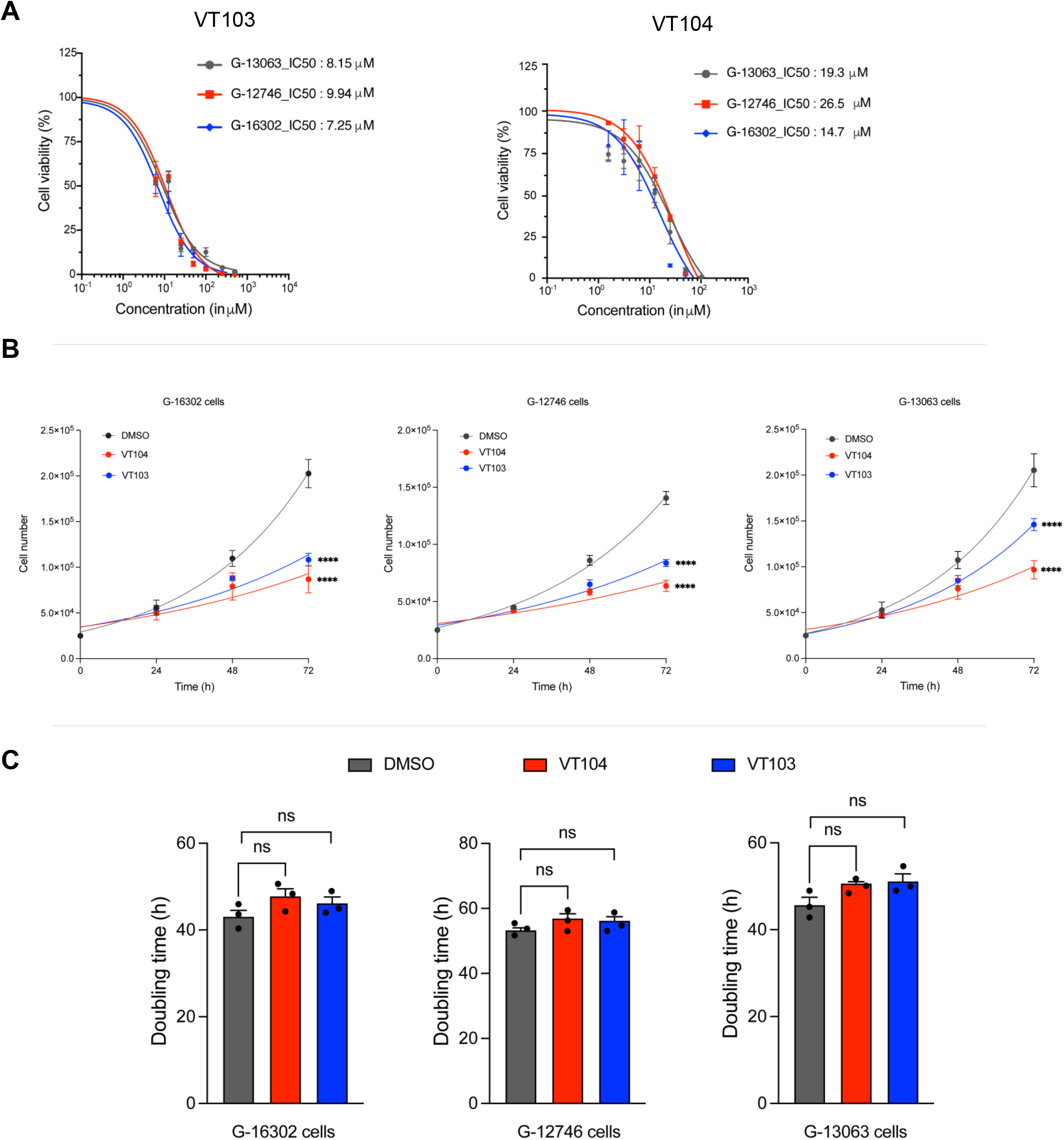
(Relates to Figure 1) Cell viability and proliferation analysis of VT103 and VT104 in patient-derived GBM cells **A.** Cell viability analysis of G-16302, G-12746, and G-13063 patient-derived GBM cell lines with diverse genomic drivers shows dose-dependent drug effects with comparable IC_50_ values for VT103 and VT104. **B.** VT103 and VT104 treatment (IC_50_ dose) shows significant decrease in cell proliferation at or after 48 hours of drug administration, across the different GBM cell lines tested. **C.** VT103 and VT104 treatment (IC_50_ dose) does not show significant decrease in GBM cell doubling time within the first 24 hours after drug administration. A, B, C: Data are presented as mean ± SEM (n=3 wells per treatment group). Statistical significance determined by two-way Student’s t-test.

**Supplementary Figure S2.**
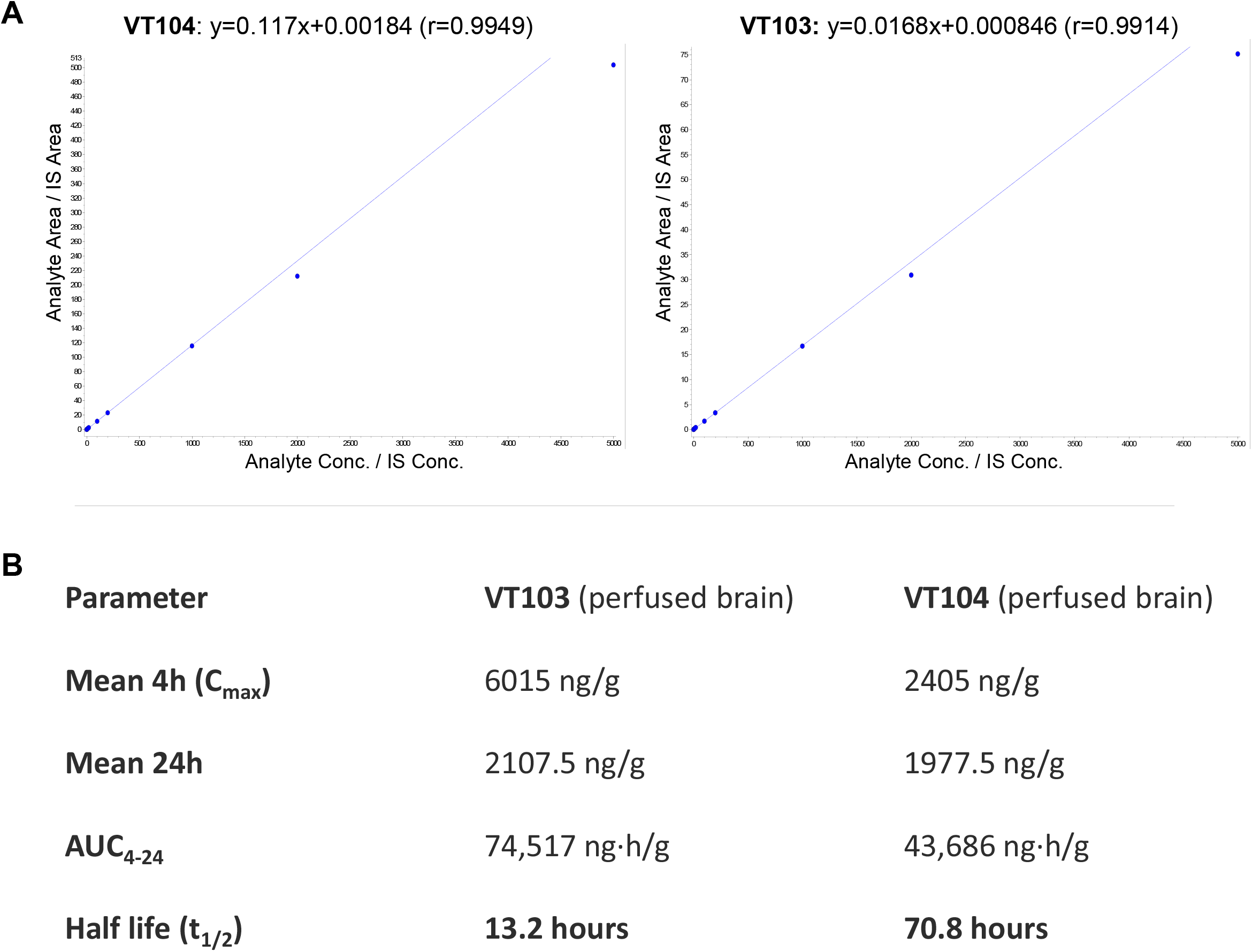
(Relates to Figure 2) Analysis of VT103 and VT104 pharmacokinetics in brain and plasma of tumor-bearing PDX mice **A.** Standard curves in mouse plasma for VT104 (left) and VT103 (right) analytes used to determine quality control and accuracy of LC-MS/MS measurements. Calibration standards and quality control samples were prepared by spiking the test compounds into blank mouse plasma and processed in parallel with the unknown samples in the same batch. **B.** Calculations of pharmacokinetics parameters for VT103 and VT104 in perfused brain tissue, including elimination half-life (t_1/2_), total drug concentration (AUC_4-24_) and maximum drug concentration (C_max_). AUC_4-24_ was calculated using the logarithmic trapezoidal method to account for the exponential nature of drug elimination. VT104 demonstrates longer drug half-life compared to VT103. VT103 and VT104 drug concentrations were measured in plasma and perfused brain tissue at 4h and 24h after last administration.

**Supplementary Figure S3.**
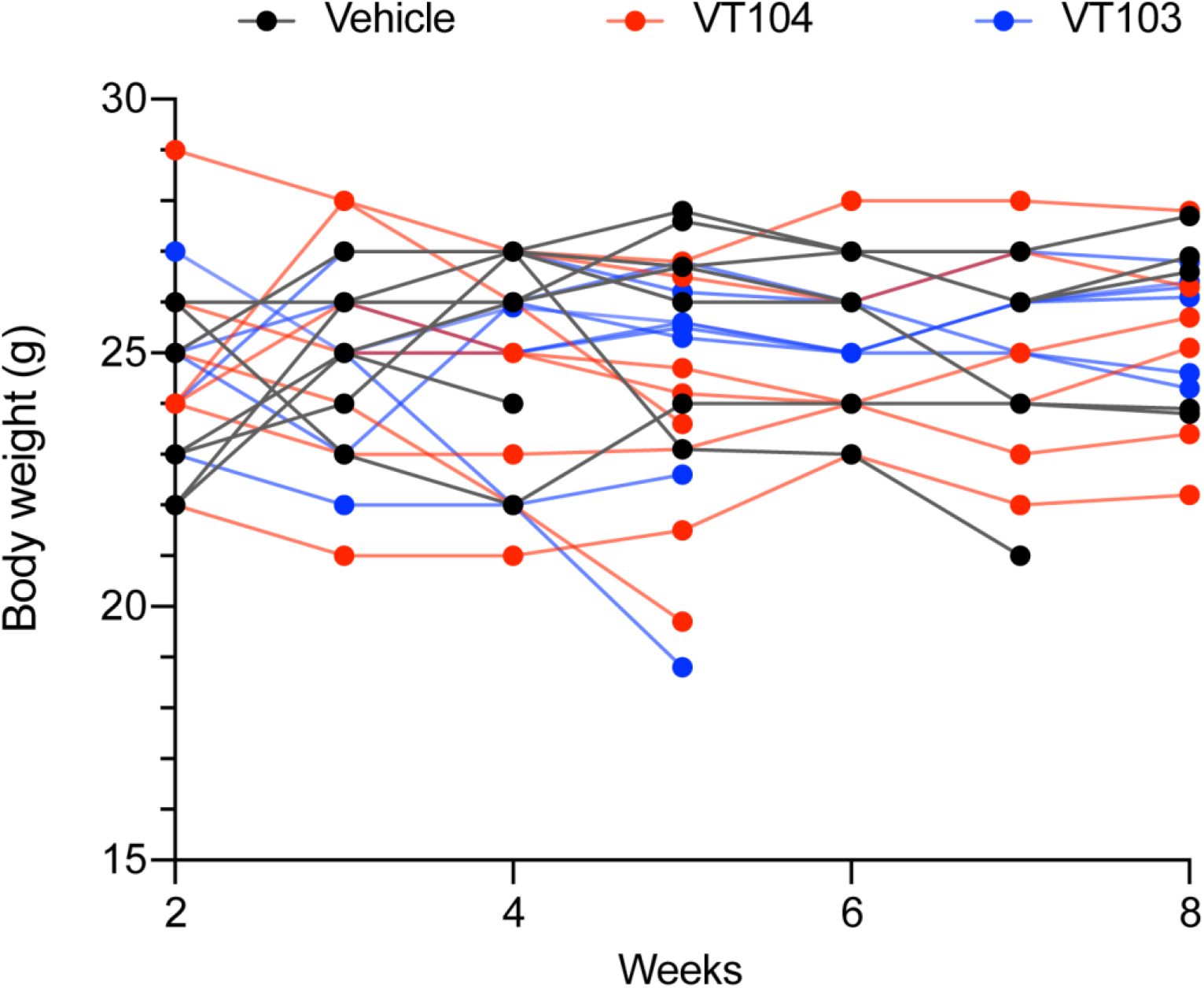
(relates to Figure 2) Body weight of G-16302 PDX mice used for pharmacokinetic studies. No substantial weight loss is seen after 7 weeks of continuous VT103 or VT104 drug administration in mice sacrificed acutely after last drug dose, which suggests a lack of systemic drug toxicity.

**Supplementary Figure S4.**
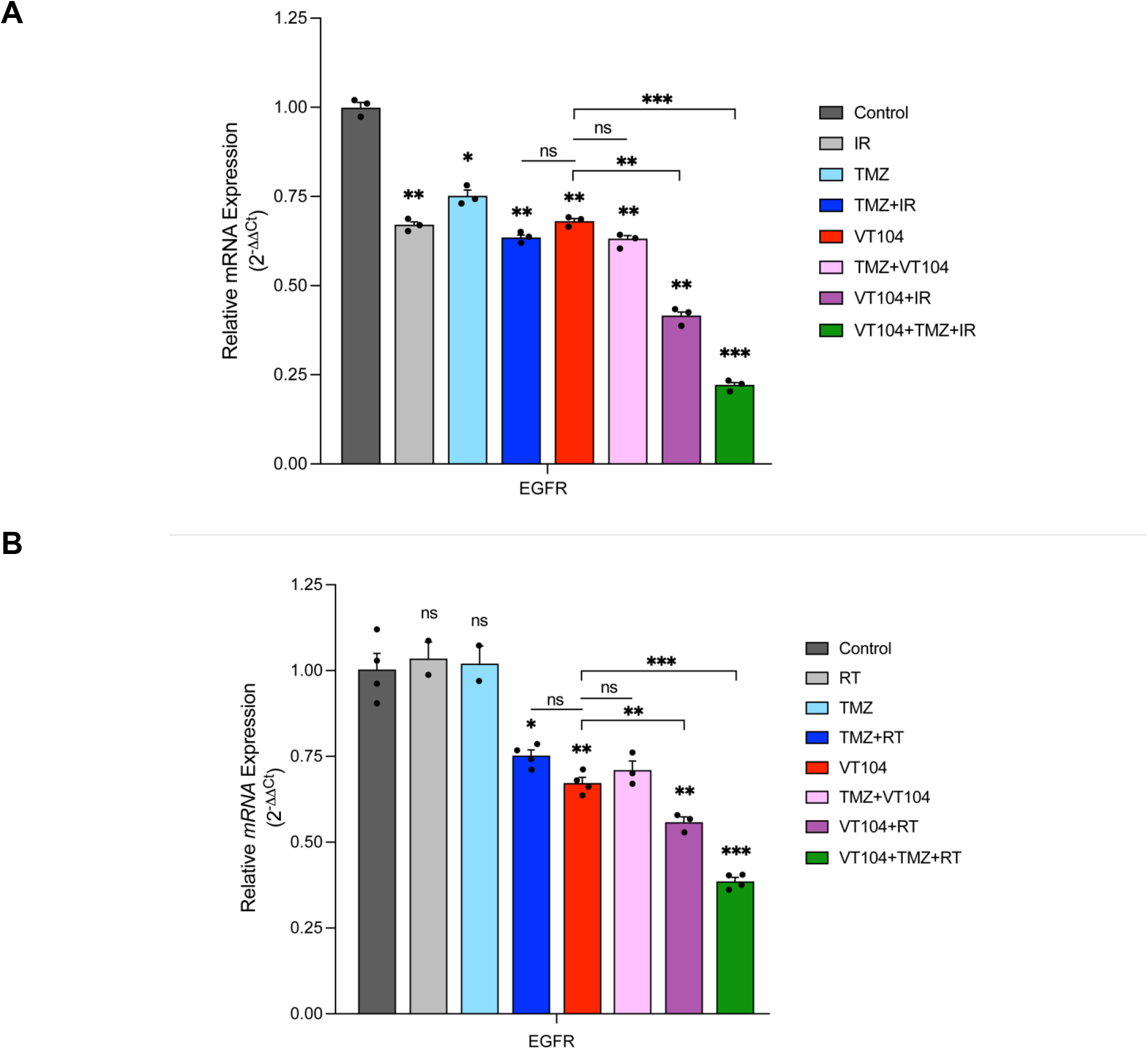
(relates to Figures 6 and 7) EGFR expression changes in patient-derived G-12315 recurrent GBM model after in-vitro and in-vivo drug combination treatments **A.** Analysis of *EGFR* expression in G-12315 PDX cells after treatment(s), showing greatest downregulation after triple combination (VT104+TMZ+RT) treatment. Bar plot represents averaged gene expression normalized to housekeeping *18S* gene. Data are presented as mean ± SEM (n=3 wells per treatment group). **B.** Analysis of *EGFR* expression in human gliomas harvested from G-12315 PDX mice, showing greatest downregulation after triple combination (VT104+TMZ+RT) treatment. Treatment began three weeks after tumor implantation (VT104: 10mg/kg/day p.o. for 3 weeks; TMZ: 25mg/kg/day p.o. for 5+5 days; RT: 2Gy/day for 5+5 days, 10Gy total). Bar plot represents averaged gene expression normalized to housekeeping *RPLPO* gene. Data are presented as mean ± SEM (n=4 mice per treatment group).

**Supplementary Figure S5.**
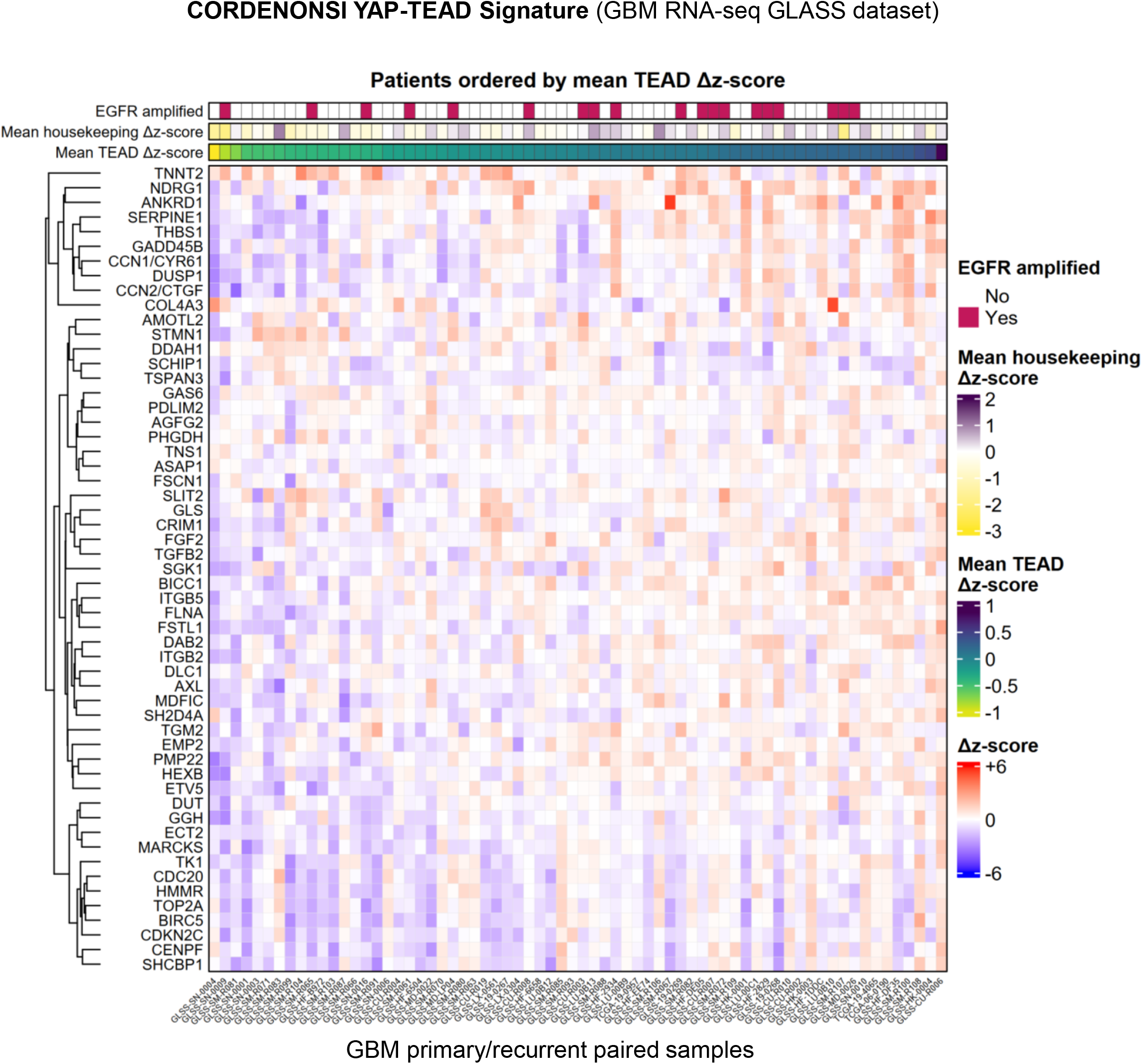
(relates to Figure 7) Longitudinal YAP–TEAD signature changes in matched primary and recurrent GBM Global analysis of TEAD-target gene RNA-seq expression in paired primary-recurrent IDH- wildtype GBM patient samples obtained from the longitudinal GLASS consortium (n=68 primary tumor samples with no treatment, n=68 patient-matched first recurrences, RT+TMZ treated). Heatmap shows recurrent minus primary mRNA z-score differences for CORDENONSI_YAP_CONSERVED_SIGNATURE genes (MSigDB) across matched GLASS glioma cases. Each column represents a patient, ordered by increasing mean YAP–TEAD signature score difference between recurrence and primary tumor. Each row represents a gene, hierarchically clustered. Red indicates higher expression in recurrence relative to matched primary tumor, blue indicates lower expression in recurrence, and white indicates no change. The heatmap scale was fixed from –6 to +6 z-score difference. The top annotations indicate patient-level EGFR amplification status and mean YAP–TEAD signature score difference.

